# Emerging Beetle-Pathogen Symbioses and Their Consequences for Forest Health: Lessons from Rapid ‘Ōhi’a Death in Hawai’i

**DOI:** 10.64898/2026.06.24.732210

**Authors:** Alexandra Boren, Sven Weber, Lisa M. Keith, Rosemary Gillespie, George Roderick, Kylle Roy

## Abstract

Invasive ambrosia beetles and fungal pathogens threaten forest ecosystems worldwide, exemplified in Hawai’i by the widespread loss of keystone species ‘ōhi’a (*Metrosideros polymorpha*), due to Rapid ‘Ōhi’a Death (ROD). A unique occurrence of five ambrosia beetle species (one native, four introduced) that vary in their symbiotic relationships with two introduced fungal pathogens provide an opportunity to test hypotheses of how opportunistic symbioses facilitate disease dynamics involving dominant forest trees. ROD is caused by two novel *Ceratocystis* fungal pathogens whose spores can spread via association with ambrosia beetles as they bore into ‘ōhi’a trees. We examined beetle-pathogen interactions of all five ambrosia beetle species in three ROD-affected regions on Hawai’i Island, and used quantitative PCR (qPCR) to provide the first molecular confirmation of the two ROD pathogens associated with the exterior, mycangia, and gut of each beetle species. Results from generalized linear models and correlation networks show that pathogen acquisition and transport, including the potential for consumption and the presence of the pathogens, are determined by beetle invasion status and mycangia morphology. A niche construction framework suggests that both varying symbioses and opportunism facilitate disease spread, with the three invasive *Xyleborus* species emerging as key disease vectors. Identifying the beetle species that are more likely to contribute to disease spread, and understanding their biology as vectors, can inform targeted conservation strategies for ‘ōhi’a and for insect-pathogen threats in forests worldwide, and illustrates the potential ecosystem-level impacts of novel and opportunistic symbioses between globally distributed invasive vectors and pathogens.

## 1. Introduction

Invasive insects and diseases are major threats to forest ecosystems (Trumbore et al. 2015). Increased globalization and climate change are contributing to the decline of native forest plant species by facilitating the invasion and spread of associated pests and pathogens (Ramsfield et al. 2016). Expanding global trade networks have created new routes for the spread of nonnative populations (Hulme 2009, Wingfield et al. 2015), and expanding ranges of many invasive forest pests associated with climate change is exacerbating their deleterious impacts on forest ecosystems (Diez et al. 2012). Island forest ecosystems, as seen in the Hawaiian Islands, are particularly vulnerable to these effects due to high endemism and the isolated evolution of many native species, leaving niches that can readily be filled by invasive populations (Russell et al. 2017). Critical to conservation of island forests is an understanding of disease dynamics, pathogens, and disease vectors.

A striking example of the vulnerability of island forest ecosystems can be found in Rapid ‘Ōhi’a Death (ROD), a novel fungal disease complex that is devastating the native tree ‘ōhi’a lehua (’ōhi’a; *Metrosideros polymorpha*) across the state of Hawai’i and whose transmission is associated with ambrosia beetles. ‘Ōhi’a are the most abundant native trees in Hawai’i (Luiz et al. 2023), and this foundational species provides habitat for many threatened and endangered species while protecting the watersheds across the state (Yelenik et al. 2020). In addition, the wood, flowers, and leaves of ‘ōhi’a have cultural importance, showcased by use in local architecture and traditional lei (Abbott 1992). ROD severely threatens Hawaiian forests, with estimates of over 1 million ‘ōhi’a trees killed by ROD since 2010 (Cannon et al. 2022).

ROD is caused by two fungal pathogens in the genus *Ceratocystis, C. lukuohia* and *C. huliohia* (Barnes et al. 2018). Both pathogens can kill ‘ōhi’a by infecting the sapwood, vascular cambium, and phloem of the tree. *C. lukuohia* belongs to the aggressive Latin American clade of *Ceratocystis* and causes more rapid and ubiquitous crown wilt, whereas *C. huliohia* is from the Asian-Australian clade of *Ceratocystis* and causes slower tree mortality by forming localized areas of necrotic tissue (Juzwik et al. 2024, Hughes et al. 2020, Barnes et al. 2018). Both pathogens can concurrently cause the same ROD outbreak, but *C. lukuohia* is considered dominant and more virulent (Barnes et al. 2018). Microsatellite analyses indicate that each represents a clonal lineage, consistent with a relatively recent introduction to Hawai’i; both were likely introduced with horticultural plants (Barnes et al. 2018).

Like many other *Ceratocystis* pathogens, the transmission of *C. lukuohia* and *C. huliohia* is associated with ambrosia beetles (Coleoptera: Curculionidae: Scolytinae) and their frass (Roy et al. 2019). Ambrosia beetles are fungus-farming beetles that have obligate nutritional relationships with their various fungal symbionts, collectively referred to as “ambrosia fungi” (Hulcr and Stelinski 2017, Vanderpool et al. 2018). The mutualism between ambrosia beetles and their symbionts is further maintained through chemical attraction: in addition to being attracted to the scent of ethanol produced by stressed and/or wounded trees (Kelsey and Westlind 2017), ambrosia beetles can be attracted to volatile organic compounds (VOCs) produced by their respective fungal symbionts (Hulcr et al. 2011). Spores of symbiotic fungi are stored in specialized structures on the beetles called mycangia, which are highly variable amongst ambrosia beetle species both in morphology and location on the body (Batra 1963, Li et al. 2018). The adult releases fungal spores from its mycangia, after which the fungi colonizes the xylem and provides the sole food source for the beetle and its larvae (De Fine Licht and Biedermann 2012, Huang et al. 2020). Ambrosia beetles can also be classified according to their mycangia type, with “small” mycangia (simple exoskeletal modification) typically hosting a more diverse array of fungal symbionts compared to “large” mycangia (complex internal structures) which are often more selective (Joseph and Keyhani 2021).

As of this writing, there are five known species of ambrosia beetles that are associated with the transmission of ROD in Hawai’i: one presumed native species *Xyleborus (X.) simillimus* Perkins, and four introduced species *Xyleborus affinis* Eichhoff, *Xyleborus perforans* (Wollaston), *Xyleborus ferrugineus* (Fabricius), and *Xyleborinus (Xi.) saxesenii* (Ratzburg) (Roy et al. 2020, 2023). With the exception of the native *X. simillimus* whose only known host is ‘ōhi’a, the ROD-associated ambrosia beetle species are all generalists (i.e., form galleries in a wide range of host trees) and invasive in Hawai’i (Seybold et al. 2016, Gohli et al. 2016, Roy et al. 2020).The *Xyleborus* species possess “small” paired preoral mycangia, whereas *Xyleborinus saxesenii* has a “large” elytral notch mycangia (Joseph and Keyhani 2021). All of the species can spread ROD through their frass, whereas the invasive beetles can serve as direct vectors of ROD by transporting the fungi, presumably on the outside of their bodies. *Ceratocystis* spores can stick to the exoskeletons of the ambrosia beetles as they disperse to bore into stressed trees to establish their galleries, therefore enabling the inoculation of uninfected ‘ōhi’a (Roy et al. 2023). It is unknown whether the one native species, *X. simillimus*, can serve as a direct vector.

The ambrosia beetles associated with ROD belong to the tribe Xyleborini, which includes many globally successful invaders that have contributed to the spread of destructive tree diseases worldwide (Demidko et al. 2021; Gohli et al. 2016). The invasive ROD vectors have transmitted other *Ceratocystis* pathogens (Saunders et al. 1967; Souza et al. 2013; Galdino et al. 2016) and, for example, have facilitated the spread of laurel wilt through cultivation of the pathogenic fungus, *Harringtonia lauricola,* within their mycangia (Harrington et al. 2008; Fraedrich et al. 2008; Carrillo et al. 2014). Although all invasive ROD-associated beetles host diverse fungal symbionts, *Xyleborus* species in particular show a heightened capacity to acquire novel fungi, likely due to their “small” paired preoral mycangia that supports more promiscuous fungal uptake (Carrillo et al. 2014, Kostovcik et al. 2015, Ibarra-Juarez et al. 2020, Biedermann 2020, Joseph and Keyhani 2021, Nuotclà et al. 2021). While ROD vectors are unlikely to cultivate ROD pathogens within their mycangia, their ability to uptake nonnative pathogenic fungi may amplify their impact as disease vectors, particularly given their broad host ranges.

Furthering our understanding of the ambrosia beetle vectors and their associations with the ROD pathogens is critical for ‘ōhi’a conservation. The recent identification of the ambrosia beetle vectors of ROD (Roy et al. 2023) has enabled more entomologically focused ROD management such as the use of repellents, but these efforts are constrained by a lack of knowledge on the extent to which the beetles are engaging with the pathogens, which could support more targeted management efforts to protect ‘ōhi’a populations and limit community-level impacts. The Niche Construction Theory (NCT), in which organisms modify environments in ways that alter selective pressures on themselves and other organisms, provides a framework to understand ambrosia beetle-*Ceratocystis* interactions and their impacts on forest ecosystems, which can then inform conservation strategies both in Hawai’i and in other forests around the world (Laland et al. 2016). In mutualisms, organisms co-construct a shared niche with potential spillover effects on the surrounding ecosystem (Laland et al. 2016, Six 2020). Applied to mutualisms between ambrosia beetles and fungal symbionts–in which fungi and beetles co-construct their niche within the tree through nutrient transport, detoxification of tree defenses, and VOC-mediated behaviors that maintain associations–NCT can help explain how the beetle-fungus symbiosis strength and flexibility correlates with the magnitude of ecosystem impacts (Ranger et al. 2018, Six 2020). The nutritional dependence of ambrosia beetles on their fungal mutualists promotes a strong symbiosis that can strongly impact their environment. The introduction of invasive ambrosia beetles and fungal pathogens creates a potential pathway for new beetle-fungus mutualisms to develop with the potential for disastrous consequences on the surrounding ecosystem (Six 2020).

We test hypotheses derived from the NCT for how the different beetle-fungus symbioses may be impacting the spread of ROD. Specifically, we use quantitative PCR (qPCR) to assess how the presence of *Ceratocystis* in the gut, mycangia, and exterior of wild-caught beetles varies depending on their invasive status and mycangia type–the first molecular assessment of the association between the two *Ceratocystis* ROD pathogens and ambrosia beetle vectors. If the beetles show internalization of *Ceratocystis* (present in the gut and/or mycangia), this would provide evidence for stronger co-niche construction with the pathogen, which could therefore contribute to larger and more durable effects on the forest ecosystem. In this case, the beetles may benefit from the fungus, either through consumption or through their ability to harbor and spread a lethal pathogen that increases the availability of stressed or dead host trees. Given the ability of the four invasive ambrosia beetles to acquire novel fungal symbionts in their mycangia and feed on diverse ambrosia fungi across host trees worldwide (Kostovcik et al. 2015, Osborn et al. 2023, Rassati et al. 2019), we expect both *Ceratocystis* pathogens to be present in their mycangia and gut of invasive beetles in addition to their exterior. However, we expect *Ceratocystis* to occur less frequently in beetles with more selective mycangia (i.e, *Xi. saxesenii*). Given the native *X. simillimus* is the only tree host specialist, we expect it to be less likely to benefit from the consumption or harboring of the novel *Ceratocystis* pathogens, and therefore expect *Ceratocystis* to be predominantly present on its exterior. Understanding vector-disease dynamics, and particularly how variation in beetle-fungus symbioses influences pathogen transmission, offers an opportunity to improve biosecurity surveillance and inform conservation-oriented management of forest pests.

## 2. Methods

### 2.1 Beetle collection

We collected ROD-associated ambrosia beetles from three regions on Hawai’i Island: ‘Ōlaʻa Forest Reserve in Hawaii’ Volcanoes National Park (OFR), Waiākea Forest Reserve (WFR), and Keaukaha Military Reserve (KMR) (Figure 1). GPS coordinates for the collection sites within each region are included in the supplementary materials (Table S1). These field sites were chosen based on reports of the presence of ROD. All beetle collection took place during June 2024.

**Figure 1.**
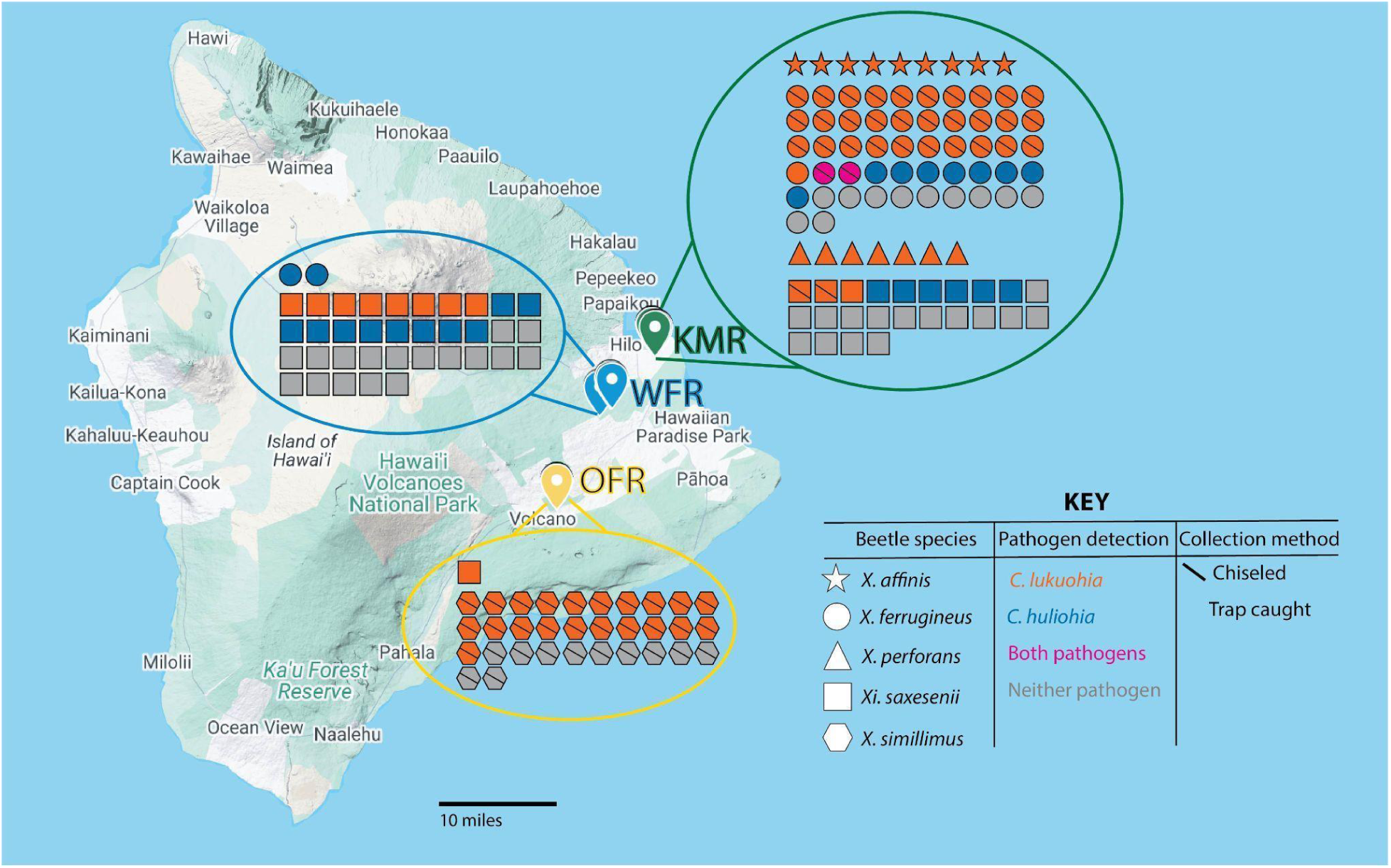
Map of beetle collection sites on Hawai’i Island. Field sites include OFR (’Ōla’a Forest Reserve in Hawai’i Volcanoes National Park), KMR (Keaukaha Military Reserve), and WFR (Waiākea Forest Reserve). The pins represent the GPS coordinates of the trap and chiseling locations; due to the proximity of the trees/traps within each site, each individual pin is not visible. Each shape represents one individual beetle that was collected; the shape dictates the species (see key). If at least one of the DNA extractions from that individual beetle (gut, mycangia, or exterior) tested positive for either of the *Ceratocystis* ROD pathogens, the individual is considered to be positive and is colored orange for the presence of *C. lukuohia*, blue for the presence of *C. huliohia*, or pink if both pathogens are present. If none of the DNA extractions from the beetle tested positive for either pathogen, the shape is colored grey. If there is a line through the shape, that beetle was chiseled from the tree. If there is no line, the beetle was caught in a Lindgren Funnel Trap. Baselayer map of Hawai’i Island sourced from Google Maps 2025.

We used 8-unit black Lindgren Funnel Traps (Contech Enterprises Inc.) to collect the four invasive beetle species (*X. affinis, X. perforans, X. ferrugineus, Xi. saxesenii*), including ten traps in WFR and ten in KMR. The traps were hung with paracord from overhanging branches of invasive vegetation ∼1-2 m above ground. Traps were placed at least 10 m apart and in close proximity to ROD-symptomatic ‘ōhi’a (i.e., the ‘ōhi’a is experiencing crown wilt and trunk staining, but retaining fine branching). Each trap was baited with a 100mL falcon tube filled with 50mL of 100% ethanol, which has worked effectively as an attractant for all four species (Roy et al. 2023). Ethanol is a by-product of disrupted aerobic respiration that is produced by stressed trees (Kelsey and Westlind 2017) and in the ROD pathosystem, the ambrosia beetles are attracted to wounded and/or stressed ‘ōhi’a trees that produce ethanol (Roy et al. 2023, Roy et al. 2025). We attached dry collection cups containing 2 crumpled Kimwipes (Kimberly-Clark Professional, Roswell, GA, USA) and a Vaportape II insecticide strip (Hercon Environmental, Emigsville, PA, USA) to the bottom of each trap. We left the traps overnight and would return to replace the trap with a new collection cup the following day. The collection cups that were removed were placed in individual gallon Ziploc (Racine, WI, USA) bags and transported to the nearby laboratory at the US Forest Service Institute of Pacific Islands Forestry for species identification. We prioritized using beetles from collection cups that were not flooded from rain to avoid cross-contamination, but when searching for species that were more difficult to find (primarily *X. affinis* and *X. perforans*) we used whichever beetles of those species we were able to find, including beetles from flooded cups. We recorded whether the cup was flooded and took that into account in our statistical models. Beetle collection metadata is available in the supplementary materials (Table S2).

Ethanol lures have not been an effective strategy for capturing the native *X. simillimus* on Hawai’i Island, likely due in part to the species’ increasingly low density, driven by competition with invasive ambrosia beetles and habitat loss (Roy et al. 2020, Roy et al. 2023). In addition, they may be attracted to host-specific kairomones emitted from ‘ōhi’a, their only known host (Samuelson 1981). Therefore, to collect *X. simillimus*, we chiseled into trees recently infected with ROD at ‘Ōlaʻa Forest Reserve (OFR). OFR is the least disturbed of our three sites and is the only site of our three in which *X. simillimus* can still consistently be found. We located active beetle galleries on the ‘ōhi’a tree by searching for gallery openings that had recently produced frass. We used a chisel and hammer to dig 1-2 cm into the wood ∼1 cm above the gallery opening. Consistent with best practices for field collection on ‘ōhi’a, all wounds made in the trees from chiseling were sealed with Spectracide Pruning Seal (Middleton, WI, USA) before leaving the site (Hughes et al. 2021, Greenplate et al. 2023). If a beetle was extracted from the resulting piece of wood, it was placed in an individual 1.5 mL tube for further processing. We additionally set up three ethanol-baited Lindgren Funnel Traps in OFR and replaced each collection cup three times over the course of two weeks. While we did not find any *X. simillimus* in the traps, we did collect one *Xi. saxesenii*. We also chiseled for *X. ferrugineus* at KMR to increase our sample size, since they are abundant at low elevation on the lower trunk of ROD-infested ‘ōhi’a (Roy et al. 2019; 2020).

The beetles were identified using morphological characteristics described by Samuelson (1981) and Wood (1982). *Xi. saxesenii* (n = 61), *X. ferrugineus* (n = 50), and *X. simillimus* (n = 32) were relatively more abundant in our chosen sample sites–consistent with prior reports of their abundance in ROD-infected ‘ōhi’a (Roy et al. 2020)–and therefore easier to collect than *X. affinis* (n = 9) or *X. perforans* (n = 7). We collected all the *X. simillimus* and roughly half of the *X. ferrugineus* (n = 32) by chiseling into ROD-positive trees; the *X. simillimus* limited to high elevation OFR, and the chiseled *X. ferrugineus* from low elevation KMR. Only *X. simillimus* and *X. ferrugineus* are able to be reliably chiseled from ‘ōhi’a because their galleries are more commonly present in the lower portion of trees.

### 2.2 Beetle processing and DNA extractions

We processed and dissected the beetles before extracting DNA from the abdomen, mycangia, and exterior wash of each beetle. We processed the beetles no later than one day after collection, storing the beetles in individual 1.5mL tubes at 4°C between identification and processing. Methods were based on modified protocols used in Roy (2023). We used forceps cleaned with 95% ethanol before being flame sterilized for transferring the beetles between tubes, and for dissection. While in the 1.5mL tube, we washed each beetle with 500uL purified water, vortexed it for ten seconds, and let the beetle sit in the water for at least ten minutes. The beetle was then moved to a new 1.5mL tube with sterile forceps, and this exterior wash was kept aside for later DNA extraction. Once in the new tube, we performed the following washes on each beetle, moving the beetle to a new 1.5mL tube in between each step: 500uL 75% ethanol; 500uL purified water; 500uL purified water. We vortexed each wash stage for ten seconds. We then dissected each beetle under a stereoscopic microscope to separate the mycangia and abdomen (Figure S1). For beetles of the *Xyleborus* genus, we separated the head (contains paired preoral mycangia) from the abdomen. For *Xyleborinus saxesenii*, we removed both elytra (contains elytral notch mycangia), separated the head from the abdomen, and discarded the head. We placed each mycangia (the head or the elytra) and each abdomen in individual 2mL tubes that each contained six 3mm zirconium grinding beads (OPS Diagnostics, Lebanon, NJ, USA) for maceration. To extract the DNA of the macerated mycangia and gut, we used the Macherey-Nagel Plant II Kit and followed the protocol for extracting genomic DNA from plants (Macherey-Nagel 2023). We used PrepMan Ultra (Thermo Fisher Scientific, Waltham, MA, USA) to extract the DNA of the exterior washes since this is a faster but similarly robust DNA extraction protocol that enabled all DNA extractions to occur promptly after beetle collection, which is necessary to maximize the quality and quantity of the extracted DNA. We followed the protocol for extracting samples for bacterial and fungal testing (Thermo Fisher Scientific, 2018) with the modification of using 50µL PrepMan Ultra with 50µL of the exterior wash for each sample. All DNA extractions were kept frozen until used in qPCR.

### 2.3 qPCR to detect *Ceratocystis* pathogens

To determine the presence of *C. lukuohia* and *C. huliohia* in the beetles, we used a high-sensitivity qPCR assay from Heller et al. (2023) on the DNA extractions from the abdomen, mycangia, and exterior wash of each beetle. This assay targets the first internal transcribed spacer (ITS) region of the multi-copy ribosomal DNA operon from the *Ceratocystis* pathogens, using probes and primer pairs designed to recognize *C. lukuohia* and *C. huliohia*, respectively (Table S3, see Heller et al. 2023). All qPCR was performed on an Applied Biosystems QuantStudio 5 instrument (Thermo Fisher, Waltham, MA). Each 20-µL reaction contained 10uL SensiFAST Probe reaction mix (Meridian Bioscience, Cincinnati, OH), 500nM primer, 200nM probe, and 5uL extracted DNA. We amplified spiked-in *Metrosideros polymorpha* DNA (10 pg/reaction) as an internal control, multiplexing the MeNu47 qPCR assay (Heller and Keith 2018) with the ITS assays (Heller et al. 2023). The thermocycling protocol consisted of a 2 minute initial denaturation at 95°C, followed by 50 cycles of 10 seconds at 95°C and 30 seconds at 60°C. Each sample was run in duplicate for both the *C. lukuohia* and *C. huliohia* assays. Samples were only re-run if they had a dubious amplification curve with an abnormally low Cq compared to the rest of the samples (<15). We also assessed the presence of *C. lukuohia* and *C. huliohia* in drill shavings from the chiseled ‘ōhi’a trees using a standard cerato-platanin extraction and qPCR protocol described in Heller and Keith (2018).

### 2.4 Categorical drivers of *Ceratocystis* presence

All analyses were performed using R (version 4.4.2; R Core Team, 2024). We used generalized linear models (GLM; glm in R) to evaluate the factors predicting the presence of the *Ceratocystis* pathogens in the body parts of each beetle species. We did not differentiate between *C. lukuohia* presence and *C. huliohia* presence in our model because the more virulent *C. lukuohia* dominated detections (see above, Figure 2), and thus counted the presence of either fungal pathogen as a positive result and the absence of both fungal pathogens as a negative result. The complete list of predictor variables we collected data on include species; body part (exterior, mycangia, gut); collection site (WFR, KFR, OFR); collection method (trap or tree chiseling); and if the beetle was collected from a flooded trap.

**Figure 2.**
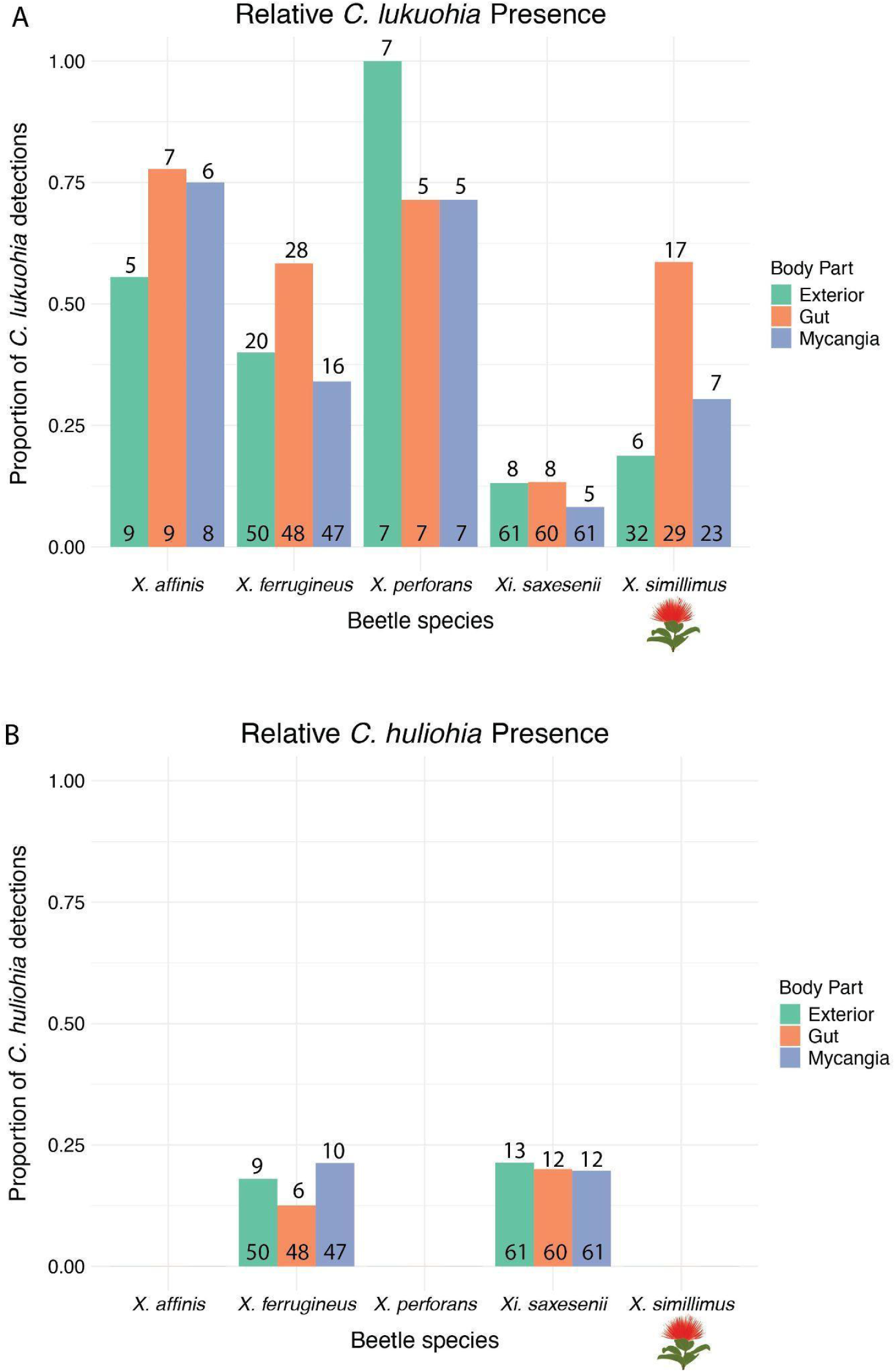
**(A)** The proportion of *C. lukuohia* detections and **(B)** the proportion of *C. huliohia* detections per body part across the ROD-associated beetle species. The values on top of each bar are the counts of *C. lukuohia* or *C. huliohia* detections, respectively, for each body part of the beetle species and the values within bars are sample sizes. The one native and specialist species (*X. simillimus*) is indicated by the ʻōhiʻa lehua in this and other figures (image courtesy of the O’ahu Invasive Species Committee).

We ran an initial GLM that included beetle species, body part, collection site, and collection method as predictor variables for the presence of *Ceratocystis* and then computed variance inflation factors (VIF) using the car package (Fox and Weisberg 2019) to assess the presence of multicollinearity between the explanatory variables. We decided to exclude the collection site from the GLM because its VIF was roughly ten-fold higher than the other explanatory variables; this indicates high collinearity, which aligns with our collection methods in which *X. simillimus* is only found at OFR. We then sequentially removed variables and compared each reduced model using analysis of deviance tests. We used the performance package (Lüdecke et al. 2021) to check all model assumptions, including the distribution of residuals, the significance of any outlying residuals, the uniformity of residuals, and the collinearity of the explanatory variables. We chose the best GLM by using the performance package to compute the Akaike Information Criteria (AIC), Tjur’s R2 for the GLM, and the conditional and marginal R2 for the GLMM as indicators of model performance. We also computed the Area Under the Curve (AUC) from the Receiver Operating Characteristic (ROC) analysis using the pROC package (Robin et al. 2011). We used the gtsummary package (Sjoberg et al. 2021) to generate a table that includes the log odds, 95% confidence intervals, and p-values for all pairwise comparisons within each predictor variable. We then used the ggeffects package (Lüdecke 2018) to perform ANOVA to assess the overall importance of each predictor variable on *Ceratocystis* presence. The predictions for each predictor variable and their 95% confidence intervals were plotted with the sjPlot package (Lüdecke 2024). We included trap flooding as an explanatory variable in a separate GLM using only the data from beetles caught in traps to evaluate whether trap flooding significantly impacts the results. We proceeded with the same assumption checks, performance evaluation, and results table generation as described previously.

We additionally used generalized linear mixed models (GLMM; lmer in R) to include the collection site as a random effect variable to account for its contribution to variation in the qPCR results while also ensuring that the collection site is not skewing the model since the variation of pathogen presence amongst collection sites is not the focus of this study. We used the same predictor variables that we used in the GLM (i.e., species, body part, collection method) as the fixed effect variables in the GLMM. We performed chi-squared tests on each beetle species to determine if there is a significant difference in *Ceratocystis* presence amongst the body parts of that species.

### 2.5 Correlation networks of *Ceratocystis* across beetle body parts

We used pairwise qPCR Cq values of each individual beetle across the whole data set to construct correlation networks to assess the likelihood of associations of the *Ceratocystis* pathogens amongst the exterior, mycangia, and gut of the beetles. The mean relative association score for each pathway (exterior and mycangia, exterior and gut, mycangia and gut) was calculated with the average association scores for each pathway for each individual beetle. If the fungus is detected in both body parts on a beetle for a given pathway (i.e., exterior and mycangia) then the association score is 1 / |Cq value for exterior - Cq value for mycangia|. If the fungus is detected only in one body part or in neither body part on a beetle then the association score is 0. If *C. lukuohia* and *C. huliohia* were both present in one sample, the average of their Cq values was used. This was only the case for three samples, all of which had similar Cq values for both pathogens. The edge width between nodes represents the strength of the fungal association between those two body parts. We created separate correlation networks for each genus to account for their different mycangia types, since *Xyleborus* beetles have a paired preoral mycangia (located near their mouth) and *Xyleborinus* beetles have an elytral notch mycangia (located at the base of their elytra). We additionally created separate correlation networks for each of the *Xyleborus* species.

## 3. Results

### 3.1 qPCR by species and by body part

Beetles from all five species tested positive for the more virulent *C. lukuohia* in their mycangia, gut, and exterior, but detections in individuals varied by species (Figure 2; Table S4). Notably, *X. ferrugineus* had the highest total number of *C. lukuohia* positives, while *X. perforans* and *X. affinis* had the greatest relative proportion of *C. lukuohia* positives when taking into account the number of beetles collected per species. Only *X. simillimus* had a significant difference in the number of positive *C. lukuohia* results per body part (χ2 df=2, p=0.0043; Table S5). *Xyleborinus saxesenii* and *X. ferrugineus* were the only species that tested positive for *C. huliohia*, and *C. huliohia* was present across all three body parts for both species (Figure 2; Table S4). With the exception of two *X. ferrugineus*, all the beetles that tested positive for *C. huliohia* were caught in traps. A higher proportion of *Xi. saxesenii* tested positive for *C. huliohia* than *C. lukuohia*, while *C. lukuohia* was more commonly present in *X. ferrugineus* than *C. huliohia*. Interestingly, co-infection of *C. lukuohia* and *C. huliohia* was only found in three samples: the gut and mycangia of the same *X. ferrugineus* individual, and the gut of a different *X. ferrugineus*. The wood from all trees that were chiseled into tested positive for *C. lukuohia* and negative for *C. huliohia*.

### 3.2 Categorical drivers of *Ceratocystis* presence

The most robust GLM for the purposes of our study included species, body part, and collection method as explanatory variables (Tjur’s R2 = 0.134; AUC = 0.703; AIC = 583.8; Table S6). Species was a significant indicator of *Ceratocystis* presence (p = 1.717e-10). Notably, the native *X. simillimus* was significantly less likely to be positive for *Ceratocystis* when compared to the invasive *Xyleborus* beetles, but there was no significant difference in the likelihood of a *Ceratocystis* positive between *X. simillimus* and *Xi. saxesenii*, the one *Xyleborinus* species (Figure 3). Body part was not a statistically significant predictor of *Ceratocystis* result (p = 0.0569), but worthy of additional study. While there were no significant pairwise differences in the likelihood of *Ceratocystis* positive results in the beetle body parts, the likelihood comparison between the gut and mycangia was the closest to having a significant result (p = 0.088) with the gut being 1.72 times more likely to be positive, whereas the likelihood comparison between the exterior and mycangia was not different (p > 0.999). Collection method was a significant predictor of *Ceratocystis* presence (p = 0.00043), with chiseled beetles being 3.13 more likely (p < 0.001) to be positive than beetles caught in a trap. This observation aligns with our expectations, since the chiseled beetles were extracted directly from ROD infected trees and are therefore more likely to be positive for *Ceratocystis* than the beetles flying into the Lindgren Funnel Traps.

**Figure 3.**
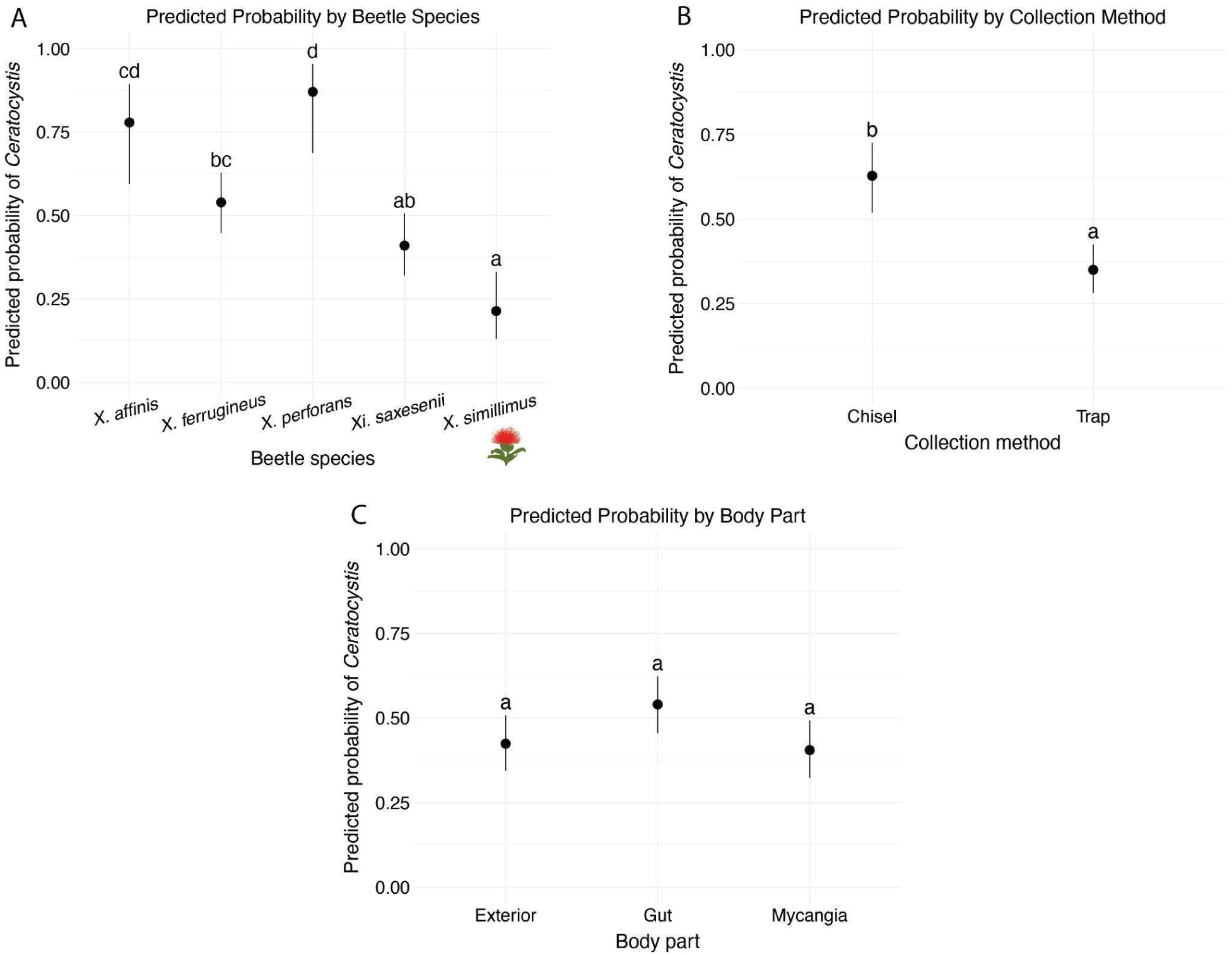
The effect of **(A)** beetle species, **(B)** collection method, and **(C)** body part on the presence of *Ceratocystis* assessed by GLM. 95% confidence intervals included. Different letters above points indicate statistically significant differences between groups (α = 0.05).

The second GLM modeled only the data from beetles caught in traps and included species, body part, and whether or not the trap was flooded as explanatory variables to assess the impact of trap flooding on *Ceratocystis* results (Tjur’s R2 = 0.122; AUC = 0.683; AIC = 357.893; Table S7). The odds ratio showed that beetles caught in dry traps were 1.78 times more likely to have a positive result for *Ceratocystis* than beetles caught in flooded traps, but this difference was not significant (p = 0.158).

Incorporating collection sites as a random effect in the GLMM created a similarly robust model compared to the GLM (AUC = 0.733; AIC = 581.9; R2(cond.) = 0.323 ; R2(marg.) = 0.267; Table S8). The most significant differences in the GLMM compared to the GLM were shown when comparing the likelihood of *Ceratocystis* positives across species. The likelihood difference between *X. simillimus* and the other *Xyleborus* species was only significant when comparing *X. simillimus* and *X. perforans*, showing that the variation between *X. simillimus* and the other *Xyleborus* species in the GLM was influenced by the collection site. The likelihoods of *Ceratocystis* positives across the pairwise comparisons of the body parts and collection methods were very similar to the results of the GLM.

### 3.3 Correlation networks show *Ceratocystis* association across beetle body parts

The correlation network showing the likelihood of *Ceratocystis* association amongst the body parts of the invasive *Xyleborus* beetles (*X. ferrugineus, X. affinis, X. perforans*) showed that the strongest likelihood of pathogen association is between the mycangia and gut (association score 1.136; Figure 4A). There was a weaker likelihood of pathogen association between the mycangia and exterior (association score 0.248) and between the exterior and gut (association score 0.356; Figure 4A). The individual correlation networks for each of the three invasive *Xyleborus* species followed a similar trend, although *X. ferrugineus* had a weaker pathogen association between its mycangia and gut when compared to *X. affinis* and *X. perforans* (Figure S2). The native *X. simillimus* had very low likelihoods of fungal association amongst all of its body parts (Figure 4C). When assessing the likelihood of *Ceratocystis* association amongst the body parts of *Xi. saxesenii*, the only *Xyleborinus* beetle, the strongest likelihood of fungal association is between the gut and exterior (association score 3.138) whereas there is a weak likelihood of fungal association between the mycangia and exterior (association score 0.640) and between the mycangia and gut (association score 0.504; Figure 4B).

**Figure 4.**
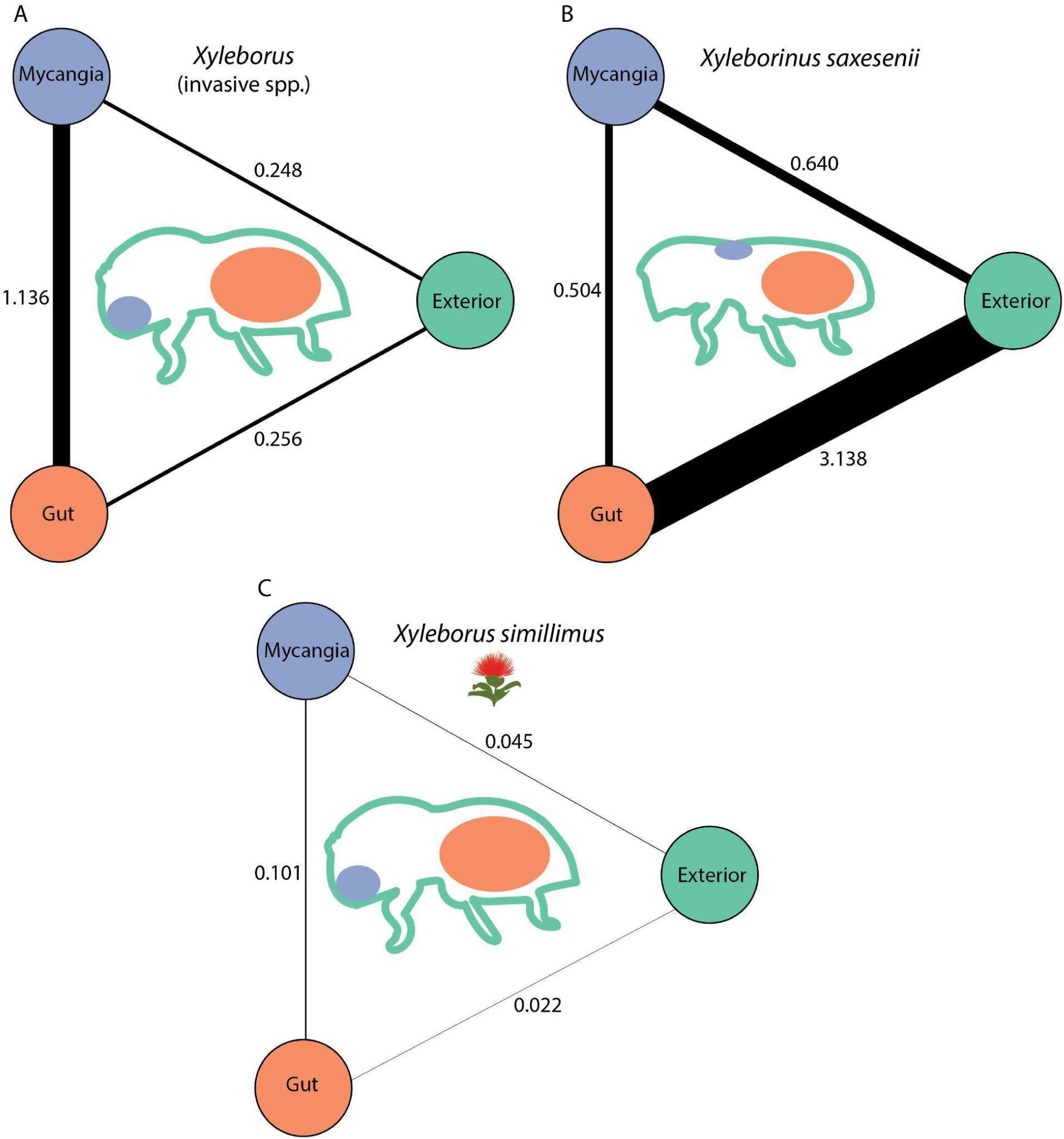
Correlation networks based on pairwise qPCR Cq values show the likelihood of *Ceratocystis* association amongst the mycangia, gut, and exterior of **(A)** the three invasive *Xyleborus* species (*X. affinis*, *X. perforans*, *X. ferrugineus*); **(B)** the one *Xyleborinus* species (*Xi. saxesenii*); and **(C)** the one native *Xyleborus* species (*X. simillimus*). The location of the mycangia for each beetle genus (*Xyleborus*–paired preoral ; *Xyleborinus*–elytral notch) is shown in purple, and the gut is shown in orange. We did not differentiate between the results for *C. lukuohia* and *C. huliohia* and instead considered the presence of either pathogen as a positive *Ceratocystis* result. Edge width directly correlates with strength of association. A higher association score indicates a stronger association.

## 4. Discussion

This study provides the first culture-independent confirmation of *Ceratocystis lukuohia* and *C. huliohia* in the mycangia, gut, and exterior across all five ROD-associated ambrosia beetle species in Hawai’i. *Ceratocystis lukuohia* was detected in all species, while *C. huliohia* was only found in *Xyleborus ferrugineus* and *Xyleborinus saxesenii*. The beetles appear to engage with *Ceratocystis* in species-specific ways, likely impacting ecosystem-level disease dynamics. These findings hold direct relevance for forest conservation in Hawai‘i, where ‘ōhi‘a (*Metrosideros polymorpha*) is a keystone species supporting watershed integrity, native biodiversity, and cultural practices. Through this assessment of beetle-pathogen associations, the study provides actionable insights for ROD management in Hawaiʻi and offers a general framework for assessing vector specificity in insect-pathogen complexes threatening global forest conservation.

### 4.1 Species-specific differences in *Ceratocystis* transmission

*Ceratocystis* prevalence varies significantly among beetle species, indicating that ROD transmission dynamics are species-specific. The invasive status, mycangia morphology, and attraction to ROD pathogens can largely explain the observed variation in pathogen presence and the recent emergence of ROD. The invasive *Xyleborus* species (*X. affinis*, *X. perforans*, and *X. ferrugineus*) had the highest overall and predicted probability of *Ceratocystis* prevalence, suggesting they are the most significant contributors to disease spread. This aligns with their paired preoral mycangia that enables promiscuous uptake of fungal symbionts and their contributions to pathogen spread globally, supporting our hypothesis that invasive *Xyleborus* exhibit stronger co-niche construction. *Xyleborinus saxesenii* exhibited the lowest proportion of pathogen presence and lacks attraction to *Ceratocystis* volatiles (Roy et al. 2025), suggesting a more passive role in transmission. Given that *X. affinis* and *X. perforans* are attracted to *C. huliohia* volatiles (Roy et al. 2025), the absence of *C. huliohia* in these species was unexpected but may reflect limited sample sizes.Despite a moderate proportion of the native *X. simillimus* carrying *C. lukuohia* DNA, its lower predicted probability of *Ceratocystis* presence, and increasing restriction to higher-elevation forests (Roy et al. 2020, Roy et al. 2023), suggest it is unlikely to significantly contribute to ROD transmission as a direct vector. Interestingly, two *X. ferrugineus* collected from the same tree harbored both *C. lukuohia* and *C. huliohia*, despite the host tree testing positive only for *C. lukuohia*. This suggests beetles may retain *Ceratocystis* from multiple trees and facilitate cross-infection, and that they may indiscriminately consume available *Ceratocystis* species.

### 4.2 Correlation networks show variable consumption of *Ceratocystis*

*Ceratocystis* was detected in the gut of all five beetle species, but correlation networks indicate that consumption differs by mycangia type and invasive status. Strong associations of *Ceratocystis* presence between mycangia and gut tissues in invasive *Xyleborus* species suggest fungal ingestion, though the proximity of their preoral mycangia to the mouth limits inference about active cultivation. In contrast, the elytral mycangia of *Xi. saxesenii* allow clearer interpretation: strong gut-exterior associations and weak mycangia-gut associations suggest passive consumption when the fungus is present in the tree rather than active cultivation. Assessing association strength among individual beetle body parts provides insight into beetle-fungus engagement beyond simple prevalence metrics and can better reflect vector capacity. For example, finding ROD in five mycangia, gut, and exterior samples across multiple beetles within one species is different from finding ROD in the mycangia, gut, and exterior of five individual beetles. The latter suggests a more sustained association, where the beetle consumes and potentially cultivates the fungus whenever it encounters it. This approach can help identify key vectors in other systems. The native *Xyleborus simillimus* exhibited weak *Ceratocystis* associations across all body parts, further supporting its limited role as a direct vector. Notably, *X. simillimus* is the only species with significantly higher *Ceratocystis* presence in its gut compared to its exterior or mycangia, likely reflecting recent consumption due to collecting all *X. simillimus* from ROD-infected ʻōhiʻa instead of from traps. The contrast between the correlation networks of invasive beetles and *X. simillimus* suggests different beetle-fungus interactions taking place.

### 4.3 Assessing symbioses between ambrosia beetle vectors and *Ceratocystis* pathogens

The origin of *C. lukuohia* and *C. huliohia* in Hawai’i remains unresolved, but understanding their ecology and evolution may explain why the native *X. simillimus* appears to interact with these pathogens differently than the invasive beetles. If either pathogen has been present in ‘ōhi’a forests for an extended period without causing disease, then a mutualistic–or at least tolerant–relationship may have developed between *Ceratocystis* and *X. simillimus*. This could offer insight into how these fungi persist in Hawaiian forests. The absence of consistent fungal associations across beetle body parts suggests *X. simillimus* may be unaffected by *Ceratocystis*, incidentally ingesting *Ceratocystis* while primarily relying on its symbiotic fungi. Further analyses of attraction to fungal volatiles or feeding behavior could clarify this relationship, and chiseling *X. simillimus* from trees with *C. huliohia* may yield different results.

The pervasiveness of *C. lukuohia* across the invasive *Xyleborus* beetles indicates it is not outcompeting symbiotic ambrosia fungi in a way that harms the beetles. Instead, the high prevalence of *Ceratocystis* in invasive beetle mycangia and gut suggests that the beetles are perhaps taking advantage of the pathogen’s presence in ROD-infected ‘ōhiʻa, either through opportunistic consumption of the fungus or through exploiting its presence to facilitate host tree death. Unlike *X. simillimus*, *Ceratocystis* may be replacing ambrosia fungi associated with the invasive beetles due to their generalist strategies and promiscuity of fungal mutualists. Our findings indicate that invasive *Xyleborus* beetles exhibit stronger co-niche construction with *Ceratocystis* than *Xi. saxesenii* or *X. simillimus*, which could lead to more durable effects of their ROD transmission in Hawaiian forests.

### 4.4 ’Ōhi’a conservation and global applications

As ROD continues to spread, informing conservation strategies is of the utmost importance to protect ‘ōhi’a. The identification of three invasive *Xyleborus* species emerging as primary vectors provides actionable insights for ROD management. Current management decisions, such as whether to apply beetle repellents to infected ‘ōhi’a, often rely on overall beetle abundance. However, if beetle abundance is low but dominated by these key *Xyleborus* species, immediate treatment may be warranted to reduce further vector establishment. Identification of primary vectors could also inform testing of biological control agents and further development of specific semiochemical attractants and repellents.

Our study provides a widely applicable framework, derived from the Niche Construction Theory and coupled with qPCR identification, for identifying primary insect vectors of forest pathogens using qPCR to inform targeted, cost-efficient management strategies that allocate their resources towards monitoring the insects with the greatest potential impacts as vectors. This approach is particularly valuable when trapping insect vectors that are attracted to oxidative stress or other pheromones, which necessitates a consideration of the species-specific variability in chemical attraction.

In addition to promoting more effective conservation strategies, these findings have bearing on ambrosia beetle dynamics beyond Hawai’i and on insect-pathogen symbioses more generally. The invasive beetles associated with ROD transmission are all globally invasive pests, and as members of the tribe Xyleborini they invade new habitats more frequently than other scolytinae beetles (Gohli et al. 2016, Rabaglia et al. 2019, Osborn et al. 2023). Most symbioses between ambrosia beetles and their fungal symbionts do not threaten natural ecosystems, but this balance can shift when beetles can spread pathogenic fungi. The invasive ambrosia beetle vectors of ROD have been associated with transmission of other fungal pathogens, but these studies have all been conducted in continental ecosystems. Assessing these symbioses in Hawai’i can clarify how island ecosystems may respond to invasive ambrosia beetles and fungal pathogens. It also further contextualizes the vector capacity of these beetles across their invasive ranges, such as how the same invasive ROD-associated beetles can also vector laurel wilt in avocado (Carillo et al. 2015).

Pathogen facilitation and opportunistic insect-pathogen symbioses are increasingly recognized as important drivers of emerging plant diseases. Co-infection and the sequence of infection can both impact disease outcomes, as seen in young grapevine decline and in the *Ascochyta* blight disease complex (May et al. 2009, Whitelaw-Weckert et al. 2013, Lamichhane and Venturi 2015). Insect vectors often intensify these dynamics by creating novel ecological partnerships with pathogens, as seen in Dutch Elm Disease transmission and in the spread of Pierce’s Disease by the introduced glassy winged sharpshooter (Roderick and Navajas 2017, Jürisoo et al. 2021). Such interactions illustrate predictions from the Niche Construction Theory, where insects and pathogens jointly modify host and ecosystem conditions in ways that reinforce their persistence and spread (Laland et al. 2016, Six 2020). As invasive ambrosia and bark beetles continue expanding their ranges, understanding these dynamics is critical, and insights from ROD directly contribute to this global framework.

The methodological implications from our molecular-based approaches also extend beyond the ROD system. qPCR is often faster, more accurate, and more sensitive than traditional phytopathogen detection methods such as culturing (McCartney et al. 2003, Schena et al. 2004, Yang and Juzwik 2017). While qPCR has been used to detect phytopathogens in some insect vectors (de Chaves et al. 2023) and to detect ambrosia fungi in ambrosia beetles (Carrillo et al. 2020), it is not commonly used to directly confirm phytopathogen presence in ambrosia or bark beetles. Our detection of *Ceratocystis* in even the smallest beetle samples (*Xi. saxesenii* elytra) highlights the utility of high-sensitivity assays (e.g., Heller et al. 2023) for other beetle-vectored phytopathogens. Wider application of qPCR to directly confirm phytopathogen presence in beetle vectors could accelerate identification of vector-pathogen associations and inform forest conservation efforts.

### 4.5 Future research on ambrosia beetle-*Ceratocystis* symbioses and ‘ōhi’a conservation

Our study provides valuable insights into details of the emerging pathogenic fungi-ambrosia beetle associations that affect ROD, but important questions persist, such as whether *Ceratocystis* integrates into the fungal microbiome or if it displaces known symbionts such as *Raffaelea* (Ophiostomatales), *Ambrosiella* (Microascales), or ascomycete yeasts (Hulcr and Stelinski 2017, Ibarra-Juarez et al. 2020, Nuotclà et al. 2021). Studies are also needed to assess the extent of *Ceratocystis* consumption during larval development, its effects on gallery symbioses, its persistence across dispersal events, and the degree of active cultivation (see De Fine Licht and Biedermann 2012). Addressing these questions will likely require culturing, RNA-based approaches, and metabarcoding (Roy et al. 2020, Wang et al. 2014).

Rapid ‘Ōhi’a Death poses a severe threat to Hawaiian forests, and our results clarify the role of ambrosia beetle species in the transmission of *Ceratocystis*, providing a foundation for targeted conservation strategies. Beyond ROD, our approach highlights the value of molecular techniques in studying insect-fungus interactions, and serves as a case study for how the Niche Construction Theory and the concept of pathogen facilitation can be applied to beetle-vectored pathogens and enhance our ability to predict and mitigate emerging forest diseases globally.

