## Supplemental Files for "Emerging Beetle-Pathogen Symbioses and Their Consequences for Forest Health: Lessons from Rapid ‘Ōhi’a Death in Hawai’i"

#### Table of Contents:

| | |
|---|---|
| **Figure S1** | Page 1 |
| **Figure S2** | Page 2 |
| **Table S1** | Page 3-4 |
| **Table S2** | Page 5-18 |
| **Table S3** | Page 19 |
| **Table S4** | Page 20 |
| **Table S5** | Page 21 |
| **Table S6** | Page 22 |
| **Table S7** | Page 23 |
| **Table S8** | Page 24 |

### Supplementary figures

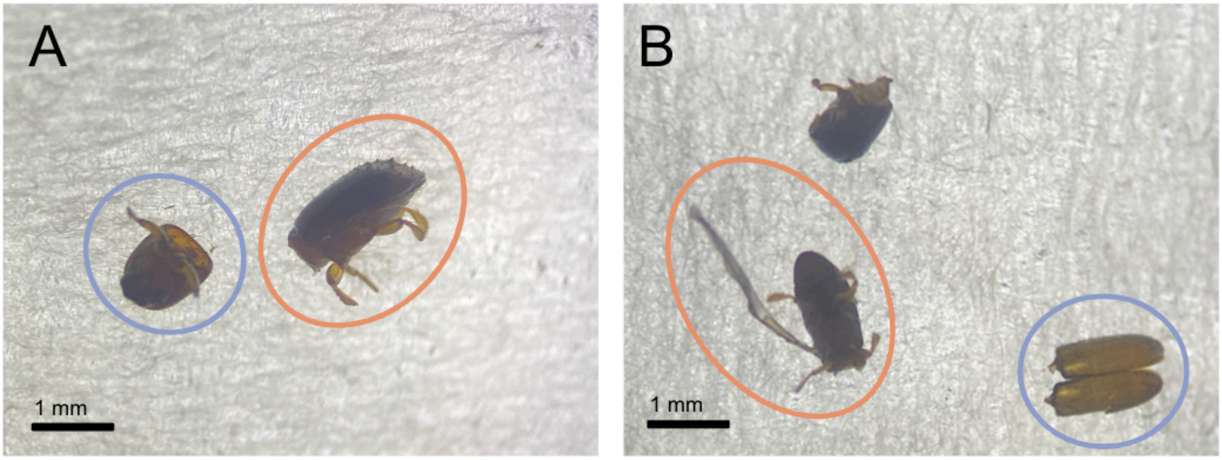

**Figure S1. Images from beetle dissection of (A) *Xyleborus* and (B) *Xyleborinus*.** The mycangia for each beetle genus (paired preoral mycangia for *Xyleborus*, and elytral notch mycangia for *Xyleborinus*) is circled in purple. The abdomen for each beetle genus is circled in orange. The representative *Xyleborus* beetle shown in (A) is *X. ferrugineus* and the beetle shown in (B) is *Xi. saxesenii*. Photos by Alexandra Boren.

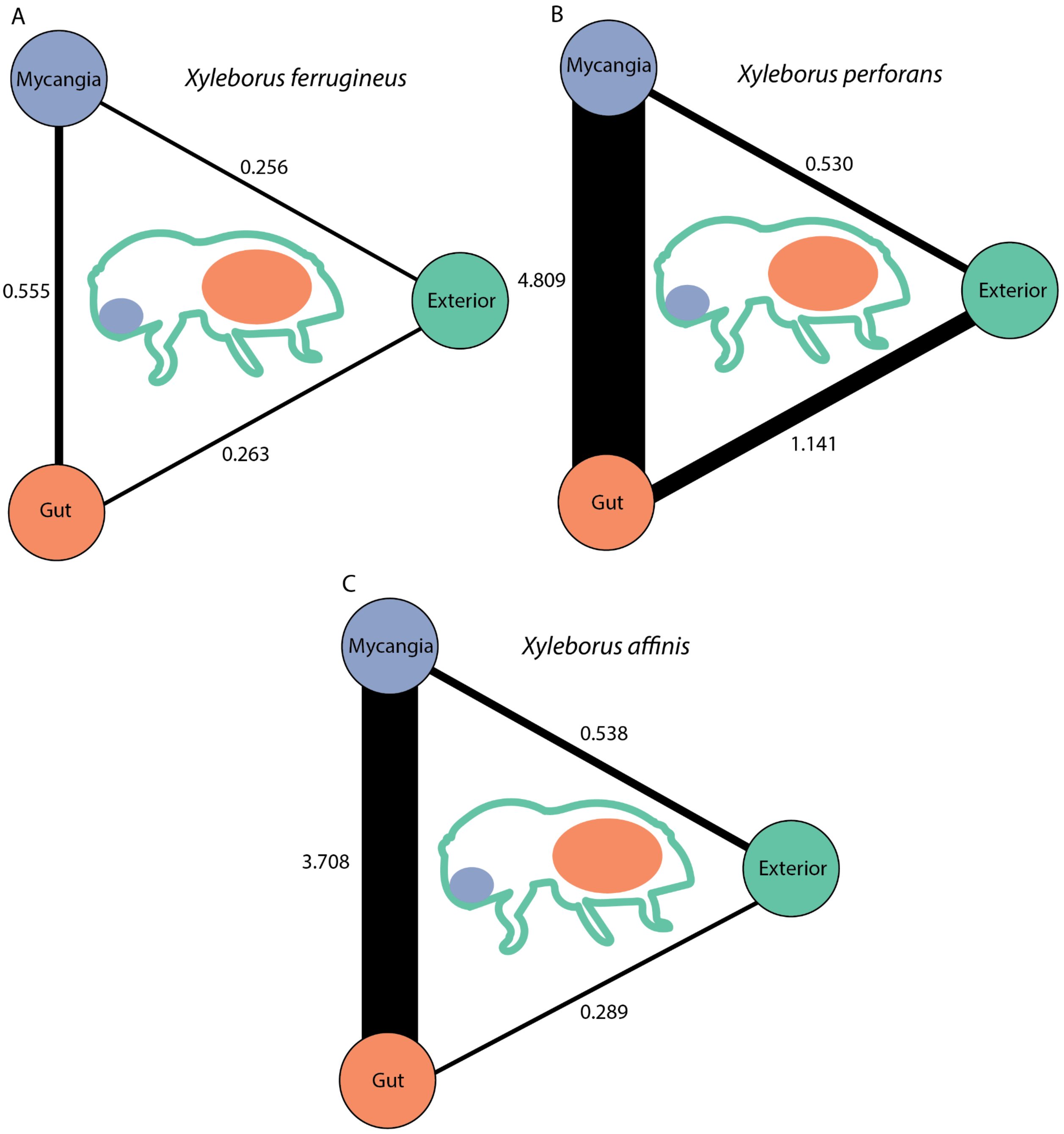

**Figure S2.** Correlation networks based on pairwise qPCR Cq values show the likelihood of *Ceratocystis* transfer amongst the mycangia, gut, and exterior of the invasive *Xyleborus* species associated with ROD: *X. affinis* **(A)**, *X. perforans* **(B)**, and *X. ferrugineus* **(C)**. We did not differentiate between the results for *C. lukuohia* and *C. huliohia* and instead considered the presence of either pathogen as a positive *Ceratocystis* result. Edge width directly correlates with strength of transfer score.

### Supplementary tables

**Table S1. GPS coordinates of beetle collection sites.** Field sites include OFR (‘Ola’a Forest Reserve in Hawai’i Volcanoes National Park), KMR (Keaukaha Military Reserve), and WFR (Waiākea Forest Reserve). The tree chiseling locations refer to the coordinates of the tree chiseled into. The Lindgren Funnel Trap locations refer to the coordinates of the traps that were hung. A total of ten traps at WFR and ten traps at KMR were used throughout the study; all traps remained in the same location for the duration of beetle collection with the exception of traps W2 and W10 which were moved to a secondary location halfway through the study (defined as W2_new and W10_new) to increase accessibility and proximity to ROD.

| **Tree chiseling locations** | | | | |
|---|---|---|---|---|
| **Site** | **Tree** | **Latitude** | **Longitude** | **Altitude** |
| OFR | OC1 | 19.443767 | -155.211808 | 1049.937622 |
| OFR | OC2 | 19.444101 | -155.211557 | 1047.49353 |
| OFR | OC3 | 19.444132 | -155.211465 | 1049.765259 |
| OFR | OC4 | 19.443883 | -155.211716 | 1021.325195 |
| KMR | KC1 | 19.708228 | -155.042122 | 54.46085 |
| KMR | KC2 | 19.708067 | -155.04227 | 51.016319 |
| KMR | KC3 | 19.708075 | -155.042292 | 51.129589 |
| KMR | KC4 | 19.708071 | -155.0423 | 51.852417 |
| KMR | KC5 | 19.708087 | -155.042332 | 49.949802 |
| KMR | KC6 | 19.708145 | -155.042335 | 50.13419 |
| KMR | KC7 | 19.708129 | -155.042332 | 49.950203 |
| KMR | KC8 | 19.70775 | -155.042169 | 51.593204 |
| **Lindgren Funnel Trap Locations** | | | | |
| **Site** | **Trap** | **Latitude** | **Longitude** | **Altitude** |
| WFR | W1 | 19.600832 | -155.135 | 508.266296 |
| WFR | W2 | 19.600972 | -155.135125 | 522.381042 |
| WFR | W2_new | 19.613409 | -155.117039 | 417.877625 |
| WFR | W3 | 19.600725 | -155.135165 | 525.66156 |
| WFR | W6 | 19.615407 | -155.114733 | 371.602386 |
| WFR | W7 | 19.615505 | -155.115134 | 418.055817 |
| WFR | W8 | 19.615446 | -155.1151 | 417.596497 |
| WFR | W09 | 19.615822 | -155.114937 | 417.355255 |
| WFR | W10 | 19.615866 | -155.114936 | 417.24057 |
| WFR | W10_new | 19.614783 | -155.114314 | 417.877626 |

| | | | | |
|---|---|---|---|---|
| KMR | K1 | 19.70218 | -155.033435 | 32.708824 |
| KMR | K2 | 19.702328 | -155.03342 | 32.639904 |
| KMR | K3 | 19.702743 | -155.033267 | 34.156029 |
| KMR | K4 | 19.702883 | -155.033271 | 33.6623 |
| KMR | K5 | 19.703072 | -155.033223 | 34.92659 |
| KMR | K6 | 19.702181 | -155.034342 | 39.693615 |
| KMR | K7 | 19.702247 | -155.034453 | 41.741146 |
| KMR | K8 | 19.702357 | -155.034634 | 43.468735 |
| KMR | K9 | 19.702392 | -155.034906 | 44.562321 |
| KMR | K10 | 19.702455 | -155.035129 | 44.806576 |
| OFR | O1 | 19.444118 | -155.211432 | 1047.284302 |
| OFR | O2 | 19.444077 | -155.211807 | 1048.464111 |
| OFR | O3 | 19.444137 | -155.211658 | 1053.783936 |

**Table S2. Collection data and qPCR results of all beetle samples.** Rows that share the same value for “Individual beetle” consist of the corresponding data from different body parts (e.g., mycangia, gut, and exterior) from the same beetle. For “Collection method”, trap refers to Lindgren Funnel Trap and chisel refers to chiseling the beetle directly from the tree. The collection sites and their abbreviations are as follows: OFR ('Ōla'a Forest Reserve in Hawai'i Volcanoes National Park), KMR (Keaukaha Military Reserve), and WFR (Waiākea Forest Reserve). A value of 1 in the columns “*C. lukuohia* presence” or “*C. huliohia* presence” indicates the presence of the respective pathogen in that sample. If the pathogen is not present (indicated by a value of 0 in “*C. lukuohia* presence” or “*C. huliohia* presence”) then the corresponding Cq average is labeled NA.

| Individual beetle | Body part | Beetle species | Collection method | Collection site | Trap flood | *C. lukuohia* presence | *C. lukuohia* Cq average | *C. huliohia* presence | *C. huliohia* Cq average |
|---|---|---|---|---|---|---|---|---|---|
| 1 | mycangia | *X. perforans* | trap | kmr | dry | 1 | 36.41 | 0 | NA |
| 1 | gut | *X. perforans* | trap | kmr | dry | 1 | 34.269 | 0 | NA |
| 1 | exterior | *X. perforans* | trap | kmr | dry | 1 | 35.154 | 0 | NA |
| 2 | mycangia | *X. perforans* | trap | kmr | dry | 1 | 38.089 | 0 | NA |
| 2 | gut | *X. perforans* | trap | kmr | dry | 0 | NA | 0 | NA |
| 2 | exterior | *X. perforans* | trap | kmr | dry | 1 | 35.158 | 0 | NA |
| 3 | mycangia | *X. perforans* | trap | kmr | dry | 0 | NA | 0 | NA |
| 3 | gut | *X. perforans* | trap | kmr | dry | 1 | 37.453 | 0 | NA |
| 3 | exterior | *X. perforans* | trap | kmr | dry | 1 | 37.236 | 0 | NA |
| 4 | mycangia | *X. perforans* | trap | kmr | dry | 0 | NA | 0 | NA |
| 4 | gut | *X. perforans* | trap | kmr | dry | 0 | NA | 0 | NA |
| 4 | exterior | *X. perforans* | trap | kmr | dry | 1 | 35.243 | 0 | NA |
| 5 | mycangia | *X. perforans* | trap | kmr | flood | 1 | 36.177 | 0 | NA |
| 5 | gut | *X. perforans* | trap | kmr | flood | 1 | 36.146 | 0 | NA |
| 5 | exterior | *X. perforans* | trap | kmr | flood | 1 | 38.037 | 0 | NA |
| 6 | mycangia | *X. perforans* | trap | kmr | flood | 1 | 34.759 | 0 | NA |
| 6 | gut | *X. perforans* | trap | kmr | flood | 1 | 36.929 | 0 | NA |
| 6 | exterior | *X. perforans* | trap | kmr | flood | 1 | 36.196 | 0 | NA |
| 7 | mycangia | *X. perforans* | trap | kmr | flood | 1 | 36.296 | 0 | NA |
| 7 | gut | *X. perforans* | trap | kmr | flood | 1 | 34.207 | 0 | NA |
| 7 | exterior | *X. perforans* | trap | kmr | flood | 1 | 37.0445 | 0 | NA |
| 8 | mycangia | *X. affinis* | trap | kmr | dry | 1 | 34.8695 | 0 | NA |
| 8 | gut | *X. affinis* | trap | kmr | dry | 0 | NA | 0 | NA |
| 8 | exterior | *X. affinis* | trap | kmr | dry | 1 | 35.898 | 0 | NA |
| 9 | gut | *X. affinis* | trap | kmr | dry | 1 | 38.404 | 0 | NA |

| | | | | | | | | | |
|---|---|---|---|---|---|---|---|---|---|
| 9 | exterior | *X. affinis* | trap | kmr | dry | 1 | 36.814 | 0 | NA |
| 10 | mycangia | *X. affinis* | trap | kmr | dry | 1 | 37.13 | 0 | NA |
| 10 | gut | *X. affinis* | trap | kmr | dry | 1 | 36.9415 | 0 | NA |
| 10 | exterior | *X. affinis* | trap | kmr | dry | 0 | NA | 0 | NA |
| 11 | mycangia | *X. affinis* | trap | kmr | dry | 0 | NA | 0 | NA |
| 11 | gut | *X. affinis* | trap | kmr | dry | 1 | 36.5515 | 0 | NA |
| 11 | exterior | *X. affinis* | trap | kmr | dry | 0 | NA | 0 | NA |
| 12 | mycangia | *X. affinis* | trap | kmr | flood | 1 | 37.058 | 0 | NA |
| 12 | gut | *X. affinis* | trap | kmr | flood | 1 | 37.0155 | 0 | NA |
| 12 | exterior | *X. affinis* | trap | kmr | flood | 1 | 37.659 | 0 | NA |
| 13 | mycangia | *X. affinis* | trap | kmr | dry | 1 | 38.005 | 0 | NA |
| 13 | gut | *X. affinis* | trap | kmr | dry | 0 | NA | 0 | NA |
| 13 | exterior | *X. affinis* | trap | kmr | dry | 1 | 37.328 | 0 | NA |
| 14 | mycangia | *X. affinis* | trap | kmr | flood | 1 | 37.4855 | 0 | NA |
| 14 | gut | *X. affinis* | trap | kmr | flood | 1 | 35.408 | 0 | NA |
| 14 | exterior | *X. affinis* | trap | kmr | flood | 0 | NA | 0 | NA |
| 15 | mycangia | *X. affinis* | trap | kmr | flood | 0 | NA | 0 | NA |
| 15 | gut | *X. affinis* | trap | kmr | flood | 1 | 36.721 | 0 | NA |
| 15 | exterior | *X. affinis* | trap | kmr | flood | 0 | NA | 0 | NA |
| 16 | mycangia | *X. affinis* | trap | kmr | flood | 1 | 34.384 | 0 | NA |
| 16 | gut | *X. affinis* | trap | kmr | flood | 1 | 37.285 | 0 | NA |
| 16 | exterior | *X. affinis* | trap | kmr | flood | 1 | 39.658 | 0 | NA |
| 17 | gut | *X. simillimus* | chisel | ofr | NA | 1 | 33.8335 | 0 | NA |
| 17 | exterior | *X. simillimus* | chisel | ofr | NA | 0 | NA | 0 | NA |
| 18 | gut | *X. simillimus* | chisel | ofr | NA | 1 | 27.649 | 0 | NA |
| 18 | exterior | *X. simillimus* | chisel | ofr | NA | 1 | 35.566 | 0 | NA |
| 19 | mycangia | *X. simillimus* | chisel | ofr | NA | 1 | 32.472 | 0 | NA |
| 19 | gut | *X. simillimus* | chisel | ofr | NA | 1 | 33.884 | 0 | NA |
| 19 | exterior | *X. simillimus* | chisel | ofr | NA | 0 | NA | 0 | NA |
| 20 | mycangia | *X. simillimus* | chisel | ofr | NA | 0 | NA | 0 | NA |
| 20 | gut | *X. simillimus* | chisel | ofr | NA | 1 | 32.9795 | 0 | NA |
| 20 | exterior | *X. simillimus* | chisel | ofr | NA | 0 | NA | 0 | NA |
| 21 | mycangia | *X. simillimus* | chisel | ofr | NA | 1 | 34.972 | 0 | NA |
| 21 | gut | *X. simillimus* | chisel | ofr | NA | 1 | 30.2375 | 0 | NA |
| 21 | exterior | *X. simillimus* | chisel | ofr | NA | 0 | NA | 0 | NA |

| | | | | | | | | | |
|---|---|---|---|---|---|---|---|---|---|
| 22 | mycangia | *X. simillimus* | chisel | ofr | NA | 1 | 34.638 | 0 | NA |
| 22 | gut | *X. simillimus* | chisel | ofr | NA | 1 | 31.924 | 0 | NA |
| 22 | exterior | *X. simillimus* | chisel | ofr | NA | 1 | 36.634 | 0 | NA |
| 23 | mycangia | *X. simillimus* | chisel | ofr | NA | 0 | NA | 0 | NA |
| 23 | gut | *X. simillimus* | chisel | ofr | NA | 1 | 34.99 | 0 | NA |
| 23 | exterior | *X. simillimus* | chisel | ofr | NA | 0 | NA | 0 | NA |
| 24 | mycangia | *X. simillimus* | chisel | ofr | NA | 0 | NA | 0 | NA |
| 24 | gut | *X. simillimus* | chisel | ofr | NA | 1 | 33.9655 | 0 | NA |
| 24 | exterior | *X. simillimus* | chisel | ofr | NA | 0 | NA | 0 | NA |
| 25 | mycangia | *X. simillimus* | chisel | ofr | NA | 0 | NA | 0 | NA |
| 25 | gut | *X. simillimus* | chisel | ofr | NA | 0 | NA | 0 | NA |
| 25 | exterior | *X. simillimus* | chisel | ofr | NA | 1 | 36.818 | 0 | NA |
| 26 | mycangia | *X. simillimus* | chisel | ofr | NA | 0 | NA | 0 | NA |
| 26 | gut | *X. simillimus* | chisel | ofr | NA | 0 | NA | 0 | NA |
| 26 | exterior | *X. simillimus* | chisel | ofr | NA | 0 | NA | 0 | NA |
| 27 | mycangia | *X. simillimus* | chisel | ofr | NA | 0 | NA | 0 | NA |
| 27 | exterior | *X. simillimus* | chisel | ofr | NA | 0 | NA | 0 | NA |
| 28 | mycangia | *X. simillimus* | chisel | ofr | NA | 0 | NA | 0 | NA |
| 28 | gut | *X. simillimus* | chisel | ofr | NA | 1 | 36.703 | 0 | NA |
| 28 | exterior | *X. simillimus* | chisel | ofr | NA | 0 | NA | 0 | NA |
| 29 | mycangia | *X. simillimus* | chisel | ofr | NA | 0 | NA | 0 | NA |
| 29 | gut | *X. simillimus* | chisel | ofr | NA | 0 | NA | 0 | NA |
| 29 | exterior | *X. simillimus* | chisel | ofr | NA | 0 | NA | 0 | NA |
| 30 | mycangia | *X. simillimus* | chisel | ofr | NA | 0 | NA | 0 | NA |
| 30 | gut | *X. simillimus* | chisel | ofr | NA | 0 | NA | 0 | NA |
| 30 | exterior | *X. simillimus* | chisel | ofr | NA | 0 | NA | 0 | NA |
| 31 | gut | *X. simillimus* | chisel | ofr | NA | 1 | 36.545 | 0 | NA |
| 31 | exterior | *X. simillimus* | chisel | ofr | NA | 0 | NA | 0 | NA |
| 32 | mycangia | *X. simillimus* | chisel | ofr | NA | 0 | NA | 0 | NA |
| 32 | gut | *X. simillimus* | chisel | ofr | NA | 1 | 37.167 | 0 | NA |
| 32 | exterior | *X. simillimus* | chisel | ofr | NA | 0 | NA | 0 | NA |
| 33 | mycangia | *X. simillimus* | chisel | ofr | NA | 0 | NA | 0 | NA |
| 33 | gut | *X. simillimus* | chisel | ofr | NA | 0 | NA | 0 | NA |
| 33 | exterior | *X. simillimus* | chisel | ofr | NA | 0 | NA | 0 | NA |
| 34 | gut | *X. simillimus* | chisel | ofr | NA | 0 | NA | 0 | NA |

| | | | | | | | | | |
|---|---|---|---|---|---|---|---|---|---|
| 34 | exterior | *X. simillimus* | chisel | ofr | NA | 0 | NA | 0 | NA |
| 35 | mycangia | *X. simillimus* | chisel | ofr | NA | 1 | 37.067 | 0 | NA |
| 35 | gut | *X. simillimus* | chisel | ofr | NA | 1 | 31.599 | 0 | NA |
| 35 | exterior | *X. simillimus* | chisel | ofr | NA | 0 | NA | 0 | NA |
| 36 | mycangia | *X. simillimus* | chisel | ofr | NA | 0 | NA | 0 | NA |
| 36 | gut | *X. simillimus* | chisel | ofr | NA | 0 | NA | 0 | NA |
| 36 | exterior | *X. simillimus* | chisel | ofr | NA | 0 | NA | 0 | NA |
| 37 | mycangia | *X. simillimus* | chisel | ofr | NA | 1 | 32.0185 | 0 | NA |
| 37 | exterior | *X. simillimus* | chisel | ofr | NA | 0 | NA | 0 | NA |
| 38 | mycangia | *X. simillimus* | chisel | ofr | NA | 1 | 36.2095 | 0 | NA |
| 38 | gut | *X. simillimus* | chisel | ofr | NA | 1 | 34.36 | 0 | NA |
| 38 | exterior | *X. simillimus* | chisel | ofr | NA | 0 | NA | 0 | NA |
| 39 | gut | *X. simillimus* | chisel | ofr | NA | 1 | 35.1315 | 0 | NA |
| 39 | exterior | *X. simillimus* | chisel | ofr | NA | 1 | 38.553 | 0 | NA |
| 40 | mycangia | *X. simillimus* | chisel | ofr | NA | 0 | NA | 0 | NA |
| 40 | gut | *X. simillimus* | chisel | ofr | NA | 0 | NA | 0 | NA |
| 40 | exterior | *X. simillimus* | chisel | ofr | NA | 0 | NA | 0 | NA |
| 41 | mycangia | *X. simillimus* | chisel | ofr | NA | 1 | 36.0775 | 0 | NA |
| 41 | exterior | *X. simillimus* | chisel | ofr | NA | 1 | 37.975 | 0 | NA |
| 42 | gut | *X. simillimus* | chisel | ofr | NA | 1 | 36.2 | 0 | NA |
| 42 | exterior | *X. simillimus* | chisel | ofr | NA | 0 | NA | 0 | NA |
| 43 | gut | *X. simillimus* | chisel | ofr | NA | 1 | 35.345 | 0 | NA |
| 43 | exterior | *X. simillimus* | chisel | ofr | NA | 0 | NA | 0 | NA |
| 44 | mycangia | *X. simillimus* | chisel | ofr | NA | 0 | NA | 0 | NA |
| 44 | gut | *X. simillimus* | chisel | ofr | NA | 1 | 29.033 | 0 | NA |
| 44 | exterior | *X. simillimus* | chisel | ofr | NA | 0 | NA | 0 | NA |
| 45 | mycangia | *X. simillimus* | chisel | ofr | NA | 0 | NA | 0 | NA |
| 45 | gut | *X. simillimus* | chisel | ofr | NA | 0 | NA | 0 | NA |
| 45 | exterior | *X. simillimus* | chisel | ofr | NA | 0 | NA | 0 | NA |
| 46 | gut | *X. simillimus* | chisel | ofr | NA | 0 | NA | 0 | NA |
| 46 | exterior | *X. simillimus* | chisel | ofr | NA | 1 | 36.716 | 0 | NA |
| 47 | gut | *X. simillimus* | chisel | ofr | NA | 0 | NA | 0 | NA |
| 47 | exterior | *X. simillimus* | chisel | ofr | NA | 0 | NA | 0 | NA |
| 48 | mycangia | *X. simillimus* | chisel | ofr | NA | 0 | NA | 0 | NA |
| 48 | gut | *X. simillimus* | chisel | ofr | NA | 0 | NA | 0 | NA |

| | | | | | | | | | |
|---|---|---|---|---|---|---|---|---|---|
| 48 | exterior | *X. simillimus* | chisel | ofr | NA | 0 | NA | 0 | NA |
| 49 | mycangia | *X. ferrugineus* | trap | kmr | dry | 0 | NA | 1 | 35.872 |
| 49 | gut | *X. ferrugineus* | trap | kmr | dry | 0 | NA | 0 | NA |
| 49 | exterior | *X. ferrugineus* | trap | kmr | dry | 0 | NA | 1 | 34.5685 |
| 50 | mycangia | *X. ferrugineus* | trap | kmr | dry | 0 | NA | 1 | 35.411 |
| 50 | gut | *X. ferrugineus* | trap | kmr | dry | 0 | NA | 0 | NA |
| 50 | exterior | *X. ferrugineus* | trap | kmr | dry | 0 | NA | 1 | 35.848 |
| 51 | mycangia | *X. ferrugineus* | trap | kmr | dry | 0 | NA | 1 | 34.803 |
| 51 | gut | *X. ferrugineus* | trap | kmr | dry | 0 | NA | 0 | NA |
| 51 | exterior | *X. ferrugineus* | trap | kmr | dry | 0 | NA | 1 | 35.568 |
| 52 | mycangia | *X. ferrugineus* | trap | kmr | dry | 0 | NA | 1 | 34.529 |
| 52 | gut | *X. ferrugineus* | trap | kmr | dry | 0 | NA | 0 | NA |
| 52 | exterior | *X. ferrugineus* | trap | kmr | dry | 0 | NA | 1 | 35.4395 |
| 53 | mycangia | *X. ferrugineus* | trap | wfr | dry | 0 | NA | 1 | 35.48 |
| 53 | gut | *X. ferrugineus* | trap | wfr | dry | 0 | NA | 1 | 37.1135 |
| 53 | exterior | *X. ferrugineus* | trap | wfr | dry | 0 | NA | 1 | 36.477 |
| 54 | mycangia | *X. ferrugineus* | trap | kmr | dry | 0 | NA | 1 | 37.689 |
| 54 | gut | *X. ferrugineus* | trap | kmr | dry | 0 | NA | 1 | 34.5185 |
| 54 | exterior | *X. ferrugineus* | trap | kmr | dry | 0 | NA | 1 | 36.136 |
| 55 | mycangia | *X. ferrugineus* | trap | wfr | dry | 0 | NA | 0 | NA |
| 55 | gut | *X. ferrugineus* | trap | wfr | dry | 0 | NA | 1 | 36.44 |
| 55 | exterior | *X. ferrugineus* | trap | wfr | dry | 0 | NA | 1 | 36.8155 |
| 56 | mycangia | *X. ferrugineus* | trap | kmr | dry | 0 | NA | 0 | NA |
| 56 | gut | *X. ferrugineus* | trap | kmr | dry | 0 | NA | 0 | NA |
| 56 | exterior | *X. ferrugineus* | trap | kmr | dry | 0 | NA | 1 | 35.566 |
| 57 | mycangia | *X. ferrugineus* | trap | kmr | dry | 0 | NA | 1 | 34.876 |
| 57 | gut | *X. ferrugineus* | trap | kmr | dry | 0 | NA | 1 | 34.718 |
| 57 | exterior | *X. ferrugineus* | trap | kmr | dry | 0 | NA | 1 | 36.316 |
| 58 | mycangia | *X. ferrugineus* | trap | kmr | dry | 0 | NA | 1 | 35.919 |
| 58 | gut | *X. ferrugineus* | trap | kmr | dry | 0 | NA | 0 | NA |
| 58 | exterior | *X. ferrugineus* | trap | kmr | dry | 0 | NA | 0 | NA |
| 59 | mycangia | *X. ferrugineus* | chisel | kmr | NA | 1 | 34.0285 | 0 | NA |
| 59 | gut | *X. ferrugineus* | chisel | kmr | NA | 1 | 35.8755 | 0 | NA |
| 59 | exterior | *X. ferrugineus* | chisel | kmr | NA | 0 | NA | 0 | NA |
| 60 | mycangia | *X. ferrugineus* | chisel | kmr | NA | 1 | 35.0115 | 1 | 34.563 |

| | | | | | | | | | |
|---|---|---|---|---|---|---|---|---|---|
| 60 | gut | *X. ferrugineus* | chisel | kmr | NA | 1 | 30.4855 | 1 | 36.3185 |
| 60 | exterior | *X. ferrugineus* | chisel | kmr | NA | 1 | 26.687 | 0 | NA |
| 61 | mycangia | *X. ferrugineus* | chisel | kmr | NA | 0 | NA | 1 | 34.8395 |
| 61 | gut | *X. ferrugineus* | chisel | kmr | NA | 1 | 33.3015 | 1 | 35.344 |
| 61 | exterior | *X. ferrugineus* | chisel | kmr | NA | 1 | 36.024 | 0 | NA |
| 62 | mycangia | *X. ferrugineus* | chisel | kmr | NA | 0 | NA | 0 | NA |
| 62 | gut | *X. ferrugineus* | chisel | kmr | NA | 1 | 33.449 | 0 | NA |
| 62 | exterior | *X. ferrugineus* | chisel | kmr | NA | 0 | NA | 0 | NA |
| 63 | mycangia | *X. ferrugineus* | chisel | kmr | NA | 1 | 35.754 | 0 | NA |
| 63 | gut | *X. ferrugineus* | chisel | kmr | NA | 0 | NA | 0 | NA |
| 63 | exterior | *X. ferrugineus* | chisel | kmr | NA | 1 | 28.538 | 0 | NA |
| 64 | mycangia | *X. ferrugineus* | chisel | kmr | NA | 0 | NA | 0 | NA |
| 64 | gut | *X. ferrugineus* | chisel | kmr | NA | 1 | 33.032 | 0 | NA |
| 64 | exterior | *X. ferrugineus* | chisel | kmr | NA | 0 | NA | 0 | NA |
| 65 | mycangia | *X. ferrugineus* | chisel | kmr | NA | 0 | NA | 0 | NA |
| 65 | gut | *X. ferrugineus* | chisel | kmr | NA | 1 | 33.0445 | 0 | NA |
| 65 | exterior | *X. ferrugineus* | chisel | kmr | NA | 1 | 28.502 | 0 | NA |
| 66 | mycangia | *X. ferrugineus* | chisel | kmr | NA | 0 | NA | 0 | NA |
| 66 | gut | *X. ferrugineus* | chisel | kmr | NA | 1 | 32.309 | 0 | NA |
| 66 | exterior | *X. ferrugineus* | chisel | kmr | NA | 0 | NA | 0 | NA |
| 67 | mycangia | *X. ferrugineus* | chisel | kmr | NA | 1 | 36.739 | 0 | NA |
| 67 | gut | *X. ferrugineus* | chisel | kmr | NA | 1 | 31.776 | 0 | NA |
| 67 | exterior | *X. ferrugineus* | chisel | kmr | NA | 1 | 29.4945 | 0 | NA |
| 68 | mycangia | *X. ferrugineus* | chisel | kmr | NA | 1 | 35.435 | 0 | NA |
| 68 | gut | *X. ferrugineus* | chisel | kmr | NA | 1 | 34.8155 | 0 | NA |
| 68 | exterior | *X. ferrugineus* | chisel | kmr | NA | 0 | NA | 0 | NA |
| 69 | mycangia | *X. ferrugineus* | chisel | kmr | NA | 1 | 33.4915 | 0 | NA |
| 69 | gut | *X. ferrugineus* | chisel | kmr | NA | 1 | 33.3615 | 0 | NA |
| 69 | exterior | *X. ferrugineus* | chisel | kmr | NA | 1 | 36.981 | 0 | NA |
| 70 | mycangia | *X. ferrugineus* | chisel | kmr | NA | 1 | 32.5615 | 0 | NA |
| 70 | gut | *X. ferrugineus* | chisel | kmr | NA | 1 | 30.212 | 0 | NA |
| 70 | exterior | *X. ferrugineus* | chisel | kmr | NA | 1 | 26.579 | 0 | NA |
| 71 | mycangia | *X. ferrugineus* | chisel | kmr | NA | 0 | NA | 0 | NA |
| 71 | gut | *X. ferrugineus* | chisel | kmr | NA | 1 | 31.6615 | 0 | NA |
| 71 | exterior | *X. ferrugineus* | chisel | kmr | NA | 1 | 30.3695 | 0 | NA |

| | | | | | | | | | |
|---|---|---|---|---|---|---|---|---|---|
| 72 | mycangia | *X. ferrugineus* | chisel | kmr | NA | 0 | NA | 0 | NA |
| 72 | gut | *X. ferrugineus* | chisel | kmr | NA | 0 | NA | 0 | NA |
| 72 | exterior | *X. ferrugineus* | chisel | kmr | NA | 1 | 37.439 | 0 | NA |
| 73 | mycangia | *X. ferrugineus* | chisel | kmr | NA | 0 | NA | 0 | NA |
| 73 | gut | *X. ferrugineus* | chisel | kmr | NA | 0 | NA | 0 | NA |
| 73 | exterior | *X. ferrugineus* | chisel | kmr | NA | 1 | 30.253 | 0 | NA |
| 74 | mycangia | *X. ferrugineus* | chisel | kmr | NA | 0 | NA | 0 | NA |
| 74 | gut | *X. ferrugineus* | chisel | kmr | NA | 1 | 36.0245 | 0 | NA |
| 74 | exterior | *X. ferrugineus* | chisel | kmr | NA | 1 | 33.475 | 0 | NA |
| 75 | mycangia | *X. ferrugineus* | chisel | kmr | NA | 0 | NA | 0 | NA |
| 75 | gut | *X. ferrugineus* | chisel | kmr | NA | 1 | 32.6615 | 0 | NA |
| 75 | exterior | *X. ferrugineus* | chisel | kmr | NA | 1 | 29.4605 | 0 | NA |
| 76 | gut | *X. ferrugineus* | chisel | kmr | NA | 1 | 31.624 | 0 | NA |
| 76 | exterior | *X. ferrugineus* | chisel | kmr | NA | 0 | NA | 0 | NA |
| 77 | mycangia | *X. ferrugineus* | chisel | kmr | NA | 1 | 32.884 | 0 | NA |
| 77 | exterior | *X. ferrugineus* | chisel | kmr | NA | 0 | NA | 0 | NA |
| 78 | mycangia | *X. ferrugineus* | chisel | kmr | NA | 1 | 35.625 | 0 | NA |
| 78 | exterior | *X. ferrugineus* | chisel | kmr | NA | 0 | NA | 0 | NA |
| 79 | gut | *X. ferrugineus* | chisel | kmr | NA | 1 | 33.9925 | 0 | NA |
| 79 | exterior | *X. ferrugineus* | chisel | kmr | NA | 0 | NA | 0 | NA |
| 80 | gut | *X. ferrugineus* | chisel | kmr | NA | 1 | 34.912 | 0 | NA |
| 80 | exterior | *X. ferrugineus* | chisel | kmr | NA | 1 | 30.044 | 0 | NA |
| 81 | mycangia | *X. ferrugineus* | chisel | kmr | NA | 0 | NA | 0 | NA |
| 81 | gut | *X. ferrugineus* | chisel | kmr | NA | 1 | 34.643 | 0 | NA |
| 81 | exterior | *X. ferrugineus* | chisel | kmr | NA | 0 | NA | 0 | NA |
| 82 | mycangia | *X. ferrugineus* | chisel | kmr | NA | 1 | 35.08 | 0 | NA |
| 82 | gut | *X. ferrugineus* | chisel | kmr | NA | 1 | 35.986 | 0 | NA |
| 82 | exterior | *X. ferrugineus* | chisel | kmr | NA | 1 | 29.681 | 0 | NA |
| 83 | mycangia | *X. ferrugineus* | chisel | kmr | NA | 1 | 34.0825 | 0 | NA |
| 83 | gut | *X. ferrugineus* | chisel | kmr | NA | 1 | 30.9545 | 0 | NA |
| 83 | exterior | *X. ferrugineus* | chisel | kmr | NA | 1 | 26.1415 | 0 | NA |
| 84 | mycangia | *X. ferrugineus* | chisel | kmr | NA | 1 | 28.8125 | 0 | NA |
| 84 | gut | *X. ferrugineus* | chisel | kmr | NA | 1 | 25.906 | 0 | NA |
| 84 | exterior | *X. ferrugineus* | chisel | kmr | NA | 1 | 34.879 | 0 | NA |
| 85 | mycangia | *X. ferrugineus* | chisel | kmr | NA | 0 | NA | 0 | NA |

| | | | | | | | | | |
|---|---|---|---|---|---|---|---|---|---|
| 85 | gut | *X. ferrugineus* | chisel | kmr | NA | 0 | NA | 0 | NA |
| 85 | exterior | *X. ferrugineus* | chisel | kmr | NA | 0 | NA | 0 | NA |
| 86 | mycangia | *X. ferrugineus* | chisel | kmr | NA | 0 | NA | 0 | NA |
| 86 | gut | *X. ferrugineus* | chisel | kmr | NA | 1 | 34.043 | 0 | NA |
| 86 | exterior | *X. ferrugineus* | chisel | kmr | NA | 0 | NA | 0 | NA |
| 87 | mycangia | *X. ferrugineus* | chisel | kmr | NA | 1 | 36.974 | 0 | NA |
| 87 | gut | *X. ferrugineus* | chisel | kmr | NA | 1 | 36.531 | 0 | NA |
| 87 | exterior | *X. ferrugineus* | chisel | kmr | NA | 1 | 32.046 | 0 | NA |
| 88 | mycangia | *X. ferrugineus* | chisel | kmr | NA | 1 | 34.1905 | 0 | NA |
| 88 | gut | *X. ferrugineus* | chisel | kmr | NA | 1 | 32.5345 | 0 | NA |
| 88 | exterior | *X. ferrugineus* | chisel | kmr | NA | 1 | 30.9465 | 0 | NA |
| 89 | mycangia | *X. ferrugineus* | chisel | kmr | NA | 1 | 35.948 | 0 | NA |
| 89 | gut | *X. ferrugineus* | chisel | kmr | NA | 1 | 33.3065 | 0 | NA |
| 89 | exterior | *X. ferrugineus* | chisel | kmr | NA | 1 | 35.482 | 0 | NA |
| 90 | mycangia | *X. ferrugineus* | trap | kmr | flood | 0 | NA | 0 | NA |
| 90 | gut | *X. ferrugineus* | trap | kmr | flood | 0 | NA | 0 | NA |
| 90 | exterior | *X. ferrugineus* | trap | kmr | flood | 0 | NA | 0 | NA |
| 91 | mycangia | *X. ferrugineus* | trap | kmr | flood | 0 | NA | 0 | NA |
| 91 | gut | *X. ferrugineus* | trap | kmr | flood | 0 | NA | 0 | NA |
| 91 | exterior | *X. ferrugineus* | trap | kmr | flood | 0 | NA | 0 | NA |
| 92 | mycangia | *X. ferrugineus* | trap | kmr | flood | 0 | NA | 0 | NA |
| 92 | gut | *X. ferrugineus* | trap | kmr | flood | 0 | NA | 0 | NA |
| 92 | exterior | *X. ferrugineus* | trap | kmr | flood | 0 | NA | 0 | NA |
| 93 | mycangia | *X. ferrugineus* | trap | kmr | dry | 0 | NA | 0 | NA |
| 93 | gut | *X. ferrugineus* | trap | kmr | dry | 1 | 38.191 | 0 | NA |
| 93 | exterior | *X. ferrugineus* | trap | kmr | dry | 0 | NA | 0 | NA |
| 94 | mycangia | *X. ferrugineus* | trap | kmr | dry | 0 | NA | 0 | NA |
| 94 | gut | *X. ferrugineus* | trap | kmr | dry | 0 | NA | 0 | NA |
| 94 | exterior | *X. ferrugineus* | trap | kmr | dry | 0 | NA | 0 | NA |
| 95 | mycangia | *X. ferrugineus* | trap | kmr | dry | 0 | NA | 0 | NA |
| 95 | gut | *X. ferrugineus* | trap | kmr | dry | 0 | NA | 0 | NA |
| 95 | exterior | *X. ferrugineus* | trap | kmr | dry | 0 | NA | 0 | NA |
| 96 | mycangia | *X. ferrugineus* | trap | kmr | flood | 0 | NA | 0 | NA |
| 96 | gut | *X. ferrugineus* | trap | kmr | flood | 0 | NA | 0 | NA |
| 96 | exterior | *X. ferrugineus* | trap | kmr | flood | 0 | NA | 0 | NA |

| | | | | | | | | | |
|---|---|---|---|---|---|---|---|---|---|
| 97 | mycangia | *X. ferrugineus* | trap | kmr | flood | 0 | NA | 0 | NA |
| 97 | gut | *X. ferrugineus* | trap | kmr | flood | 1 | 36.032 | 0 | NA |
| 97 | exterior | *X. ferrugineus* | trap | kmr | flood | 1 | 36.6115 | 0 | NA |
| 98 | mycangia | *X. ferrugineus* | trap | kmr | flood | 1 | 36.594 | 0 | NA |
| 98 | gut | *X. ferrugineus* | trap | kmr | flood | 1 | 37.656 | 0 | NA |
| 98 | exterior | *X. ferrugineus* | trap | kmr | flood | 0 | NA | 0 | NA |
| 99 | mycangia | *Xi. saxesenii* | trap | kmr | dry | 0 | NA | 0 | NA |
| 99 | gut | *Xi. saxesenii* | trap | kmr | dry | 0 | NA | 0 | NA |
| 99 | exterior | *Xi. saxesenii* | trap | kmr | dry | 0 | NA | 0 | NA |
| 100 | mycangia | *Xi. saxesenii* | trap | kmr | dry | 0 | NA | 0 | NA |
| 100 | gut | *Xi. saxesenii* | trap | kmr | dry | 0 | NA | 0 | NA |
| 100 | exterior | *Xi. saxesenii* | trap | kmr | dry | 0 | NA | 0 | NA |
| 101 | mycangia | *Xi. saxesenii* | trap | kmr | dry | 0 | NA | 0 | NA |
| 101 | gut | *Xi. saxesenii* | trap | kmr | dry | 0 | NA | 0 | NA |
| 101 | exterior | *Xi. saxesenii* | trap | kmr | dry | 0 | NA | 0 | NA |
| 102 | mycangia | *Xi. saxesenii* | trap | kmr | dry | 0 | NA | 0 | NA |
| 102 | gut | *Xi. saxesenii* | trap | kmr | dry | 0 | NA | 0 | NA |
| 102 | exterior | *Xi. saxesenii* | trap | kmr | dry | 0 | NA | 0 | NA |
| 103 | mycangia | *Xi. saxesenii* | trap | kmr | dry | 0 | NA | 0 | NA |
| 103 | gut | *Xi. saxesenii* | trap | kmr | dry | 0 | NA | 0 | NA |
| 103 | exterior | *Xi. saxesenii* | trap | kmr | dry | 0 | NA | 0 | NA |
| 104 | mycangia | *Xi. saxesenii* | trap | kmr | dry | 0 | NA | 0 | NA |
| 104 | gut | *Xi. saxesenii* | trap | kmr | dry | 0 | NA | 0 | NA |
| 104 | exterior | *Xi. saxesenii* | trap | kmr | dry | 0 | NA | 0 | NA |
| 105 | mycangia | *Xi. saxesenii* | trap | kmr | dry | 0 | NA | 0 | NA |
| 105 | gut | *Xi. saxesenii* | trap | kmr | dry | 0 | NA | 0 | NA |
| 105 | exterior | *Xi. saxesenii* | trap | kmr | dry | 0 | NA | 0 | NA |
| 106 | mycangia | *Xi. saxesenii* | trap | kmr | dry | 0 | NA | 0 | NA |
| 106 | gut | *Xi. saxesenii* | trap | kmr | dry | 0 | NA | 0 | NA |
| 106 | exterior | *Xi. saxesenii* | trap | kmr | dry | 0 | NA | 0 | NA |
| 107 | mycangia | *Xi. saxesenii* | trap | kmr | dry | 0 | NA | 0 | NA |
| 107 | gut | *Xi. saxesenii* | trap | kmr | dry | 0 | NA | 0 | NA |
| 107 | exterior | *Xi. saxesenii* | trap | kmr | dry | 0 | NA | 0 | NA |
| 108 | mycangia | *Xi. saxesenii* | trap | kmr | dry | 0 | NA | 0 | NA |
| 108 | gut | *Xi. saxesenii* | trap | kmr | dry | 0 | NA | 0 | NA |

| | | | | | | | | | |
|---|---|---|---|---|---|---|---|---|---|
| 108 | exterior | *Xi. saxesenii* | trap | kmr | dry | 0 | NA | 0 | NA |
| 109 | mycangia | *Xi. saxesenii* | trap | kmr | dry | 0 | NA | 0 | NA |
| 109 | gut | *Xi. saxesenii* | trap | kmr | dry | 0 | NA | 0 | NA |
| 109 | exterior | *Xi. saxesenii* | trap | kmr | dry | 0 | NA | 0 | NA |
| 110 | mycangia | *Xi. saxesenii* | trap | kmr | dry | 1 | 36.579 | 0 | NA |
| 110 | gut | *Xi. saxesenii* | trap | kmr | dry | 0 | NA | 0 | NA |
| 110 | exterior | *Xi. saxesenii* | trap | kmr | dry | 0 | NA | 0 | NA |
| 111 | mycangia | *Xi. saxesenii* | trap | kmr | dry | 0 | NA | 0 | NA |
| 111 | gut | *Xi. saxesenii* | trap | kmr | dry | 0 | NA | 0 | NA |
| 111 | exterior | *Xi. saxesenii* | trap | kmr | dry | 0 | NA | 0 | NA |
| 112 | mycangia | *Xi. saxesenii* | trap | kmr | dry | 0 | NA | 0 | NA |
| 112 | gut | *Xi. saxesenii* | trap | kmr | dry | 0 | NA | 0 | NA |
| 112 | exterior | *Xi. saxesenii* | trap | kmr | dry | 0 | NA | 0 | NA |
| 113 | mycangia | *Xi. saxesenii* | trap | wfr | dry | 0 | NA | 0 | NA |
| 113 | gut | *Xi. saxesenii* | trap | wfr | dry | 1 | 39.898 | 0 | NA |
| 113 | exterior | *Xi. saxesenii* | trap | wfr | dry | 0 | NA | 0 | NA |
| 114 | mycangia | *Xi. saxesenii* | trap | wfr | dry | 0 | NA | 0 | NA |
| 114 | gut | *Xi. saxesenii* | trap | wfr | dry | 0 | NA | 0 | NA |
| 114 | exterior | *Xi. saxesenii* | trap | wfr | dry | 1 | 36.3165 | 0 | NA |
| 115 | mycangia | *Xi. saxesenii* | trap | wfr | dry | 0 | NA | 0 | NA |
| 115 | gut | *Xi. saxesenii* | trap | wfr | dry | 0 | NA | 0 | NA |
| 115 | exterior | *Xi. saxesenii* | trap | wfr | dry | 0 | NA | 0 | NA |
| 116 | mycangia | *Xi. saxesenii* | trap | wfr | dry | 0 | NA | 0 | NA |
| 116 | exterior | *Xi. saxesenii* | trap | wfr | dry | 0 | NA | 0 | NA |
| 117 | mycangia | *Xi. saxesenii* | trap | wfr | dry | 0 | NA | 0 | NA |
| 117 | gut | *Xi. saxesenii* | trap | wfr | dry | 0 | NA | 0 | NA |
| 117 | exterior | *Xi. saxesenii* | trap | wfr | dry | 0 | NA | 0 | NA |
| 118 | mycangia | *Xi. saxesenii* | trap | wfr | dry | 0 | NA | 0 | NA |
| 118 | gut | *Xi. saxesenii* | trap | wfr | dry | 0 | NA | 0 | NA |
| 118 | exterior | *Xi. saxesenii* | trap | wfr | dry | 0 | NA | 0 | NA |
| 119 | mycangia | *Xi. saxesenii* | trap | wfr | dry | 0 | NA | 0 | NA |
| 119 | gut | *Xi. saxesenii* | trap | wfr | dry | 0 | NA | 0 | NA |
| 119 | exterior | *Xi. saxesenii* | trap | wfr | dry | 0 | NA | 0 | NA |
| 120 | mycangia | *Xi. saxesenii* | trap | wfr | dry | 0 | NA | 0 | NA |
| 120 | gut | *Xi. saxesenii* | trap | wfr | dry | 0 | NA | 0 | NA |

| | | | | | | | | | |
|---|---|---|---|---|---|---|---|---|---|
| 120 | exterior | *Xi. saxesenii* | trap | wfr | dry | 0 | NA | 0 | NA |
| 121 | mycangia | *Xi. saxesenii* | trap | wfr | dry | 0 | NA | 0 | NA |
| 121 | gut | *Xi. saxesenii* | trap | wfr | dry | 0 | NA | 0 | NA |
| 121 | exterior | *Xi. saxesenii* | trap | wfr | dry | 0 | NA | 0 | NA |
| 122 | mycangia | *Xi. saxesenii* | trap | wfr | dry | 0 | NA | 0 | NA |
| 122 | gut | *Xi. saxesenii* | trap | wfr | dry | 0 | NA | 0 | NA |
| 122 | exterior | *Xi. saxesenii* | trap | wfr | dry | 0 | NA | 0 | NA |
| 123 | mycangia | *Xi. saxesenii* | trap | wfr | dry | 0 | NA | 0 | NA |
| 123 | gut | *Xi. saxesenii* | trap | wfr | dry | 0 | NA | 0 | NA |
| 123 | exterior | *Xi. saxesenii* | trap | wfr | dry | 0 | NA | 0 | NA |
| 124 | mycangia | *Xi. saxesenii* | trap | wfr | dry | 0 | NA | 0 | NA |
| 124 | gut | *Xi. saxesenii* | trap | wfr | dry | 0 | NA | 0 | NA |
| 124 | exterior | *Xi. saxesenii* | trap | wfr | dry | 0 | NA | 0 | NA |
| 125 | mycangia | *Xi. saxesenii* | trap | wfr | dry | 0 | NA | 0 | NA |
| 125 | gut | *Xi. saxesenii* | trap | wfr | dry | 0 | NA | 0 | NA |
| 125 | exterior | *Xi. saxesenii* | trap | wfr | dry | 0 | NA | 0 | NA |
| 126 | mycangia | *Xi. saxesenii* | trap | wfr | dry | 0 | NA | 0 | NA |
| 126 | gut | *Xi. saxesenii* | trap | wfr | dry | 0 | NA | 0 | NA |
| 126 | exterior | *Xi. saxesenii* | trap | wfr | dry | 0 | NA | 0 | NA |
| 127 | mycangia | *Xi. saxesenii* | trap | wfr | dry | 0 | NA | 0 | NA |
| 127 | gut | *Xi. saxesenii* | trap | wfr | dry | 0 | NA | 0 | NA |
| 127 | exterior | *Xi. saxesenii* | trap | wfr | dry | 0 | NA | 0 | NA |
| 128 | mycangia | *Xi. saxesenii* | trap | wfr | dry | 0 | NA | 0 | NA |
| 128 | gut | *Xi. saxesenii* | trap | wfr | dry | 0 | NA | 0 | NA |
| 128 | exterior | *Xi. saxesenii* | trap | wfr | dry | 0 | NA | 0 | NA |
| 129 | mycangia | *Xi. saxesenii* | trap | wfr | dry | 0 | NA | 0 | NA |
| 129 | gut | *Xi. saxesenii* | trap | wfr | dry | 0 | NA | 0 | NA |
| 129 | exterior | *Xi. saxesenii* | trap | wfr | dry | 0 | NA | 0 | NA |
| 130 | mycangia | *Xi. saxesenii* | trap | wfr | dry | 0 | NA | 1 | 34.985 |
| 130 | gut | *Xi. saxesenii* | trap | wfr | dry | 0 | NA | 0 | NA |
| 130 | exterior | *Xi. saxesenii* | trap | wfr | dry | 0 | NA | 1 | 34.561 |
| 131 | mycangia | *Xi. saxesenii* | trap | wfr | dry | 0 | NA | 1 | 35.447 |
| 131 | gut | *Xi. saxesenii* | trap | wfr | dry | 0 | NA | 1 | 36.63 |
| 131 | exterior | *Xi. saxesenii* | trap | wfr | dry | 0 | NA | 1 | 35.5775 |
| 132 | mycangia | *Xi. saxesenii* | trap | wfr | dry | 0 | NA | 0 | NA |

| | | | | | | | | | |
|---|---|---|---|---|---|---|---|---|---|
| 132 | gut | *Xi. saxesenii* | trap | wfr | dry | 0 | NA | 1 | 35.32 |
| 132 | exterior | *Xi. saxesenii* | trap | wfr | dry | 0 | NA | 1 | 36.124 |
| 133 | mycangia | *Xi. saxesenii* | trap | wfr | dry | 0 | NA | 1 | 36.04 |
| 133 | gut | *Xi. saxesenii* | trap | wfr | dry | 0 | NA | 1 | 35.766 |
| 133 | exterior | *Xi. saxesenii* | trap | wfr | dry | 0 | NA | 0 | NA |
| 134 | mycangia | *Xi. saxesenii* | trap | wfr | dry | 0 | NA | 1 | 37.177 |
| 134 | gut | *Xi. saxesenii* | trap | wfr | dry | 0 | NA | 1 | 36.032 |
| 134 | exterior | *Xi. saxesenii* | trap | wfr | dry | 0 | NA | 1 | 36.305 |
| 135 | mycangia | *Xi. saxesenii* | trap | wfr | dry | 0 | NA | 1 | 36.946 |
| 135 | gut | *Xi. saxesenii* | trap | wfr | dry | 0 | NA | 1 | 33.633 |
| 135 | exterior | *Xi. saxesenii* | trap | wfr | dry | 0 | NA | 1 | 36.3045 |
| 136 | mycangia | *Xi. saxesenii* | trap | wfr | dry | 0 | NA | 1 | 35.0215 |
| 136 | gut | *Xi. saxesenii* | trap | wfr | dry | 0 | NA | 0 | NA |
| 136 | exterior | *Xi. saxesenii* | trap | wfr | dry | 0 | NA | 1 | 34.8105 |
| 137 | mycangia | *Xi. saxesenii* | trap | wfr | dry | 0 | NA | 1 | 35.857 |
| 137 | gut | *Xi. saxesenii* | trap | wfr | dry | 0 | NA | 1 | 35.114 |
| 137 | exterior | *Xi. saxesenii* | trap | wfr | dry | 0 | NA | 1 | 36.086 |
| 138 | mycangia | *Xi. saxesenii* | trap | wfr | dry | 0 | NA | 1 | 35.179 |
| 138 | gut | *Xi. saxesenii* | trap | wfr | dry | 0 | NA | 1 | 36.4315 |
| 138 | exterior | *Xi. saxesenii* | trap | wfr | dry | 0 | NA | 1 | 35.981 |
| 139 | mycangia | *Xi. saxesenii* | trap | wfr | dry | 0 | NA | 0 | NA |
| 139 | gut | *Xi. saxesenii* | trap | wfr | dry | 0 | NA | 0 | NA |
| 139 | exterior | *Xi. saxesenii* | trap | wfr | dry | 0 | NA | 0 | NA |
| 140 | mycangia | *Xi. saxesenii* | trap | wfr | dry | 0 | NA | 1 | 35.419 |
| 140 | gut | *Xi. saxesenii* | trap | wfr | dry | 0 | NA | 1 | 35.676 |
| 140 | exterior | *Xi. saxesenii* | trap | wfr | dry | 0 | NA | 1 | 36.433 |
| 141 | mycangia | *Xi. saxesenii* | trap | wfr | dry | 0 | NA | 0 | NA |
| 141 | gut | *Xi. saxesenii* | trap | wfr | dry | 0 | NA | 1 | 35.823 |
| 141 | exterior | *Xi. saxesenii* | trap | wfr | dry | 0 | NA | 1 | 36.81 |
| 142 | mycangia | *Xi. saxesenii* | trap | wfr | dry | 1 | 38.026 | 0 | NA |
| 142 | gut | *Xi. saxesenii* | trap | wfr | dry | 1 | 37.421 | 0 | NA |
| 142 | exterior | *Xi. saxesenii* | trap | wfr | dry | 1 | 37.415 | 0 | NA |
| 143 | mycangia | *Xi. saxesenii* | trap | wfr | dry | 0 | NA | 0 | NA |
| 143 | gut | *Xi. saxesenii* | trap | wfr | dry | 0 | NA | 0 | NA |
| 143 | exterior | *Xi. saxesenii* | trap | wfr | dry | 1 | 37.0285 | 0 | NA |

| | | | | | | | | | |
|---|---|---|---|---|---|---|---|---|---|
| 144 | mycangia | *Xi. saxesenii* | trap | wfr | dry | 0 | NA | 0 | NA |
| 144 | gut | *Xi. saxesenii* | trap | wfr | dry | 1 | 39.254 | 0 | NA |
| 144 | exterior | *Xi. saxesenii* | trap | wfr | dry | 0 | NA | 0 | NA |
| 145 | mycangia | *Xi. saxesenii* | trap | wfr | dry | 0 | NA | 0 | NA |
| 145 | gut | *Xi. saxesenii* | trap | wfr | dry | 1 | 36.6495 | 0 | NA |
| 145 | exterior | *Xi. saxesenii* | trap | wfr | dry | 0 | NA | 0 | NA |
| 146 | mycangia | *Xi. saxesenii* | trap | wfr | dry | 0 | NA | 0 | NA |
| 146 | gut | *Xi. saxesenii* | trap | wfr | dry | 0 | NA | 0 | NA |
| 146 | exterior | *Xi. saxesenii* | trap | wfr | dry | 0 | NA | 0 | NA |
| 147 | mycangia | *Xi. saxesenii* | trap | wfr | dry | 0 | NA | 0 | NA |
| 147 | gut | *Xi. saxesenii* | trap | wfr | dry | 0 | NA | 0 | NA |
| 147 | exterior | *Xi. saxesenii* | trap | wfr | dry | 1 | 35.062 | 0 | NA |
| 148 | mycangia | *Xi. saxesenii* | trap | wfr | dry | 1 | 36.428 | 0 | NA |
| 148 | gut | *Xi. saxesenii* | trap | wfr | dry | 1 | 36.874 | 0 | NA |
| 148 | exterior | *Xi. saxesenii* | trap | wfr | dry | 1 | 36.332 | 0 | NA |
| 149 | mycangia | *Xi. saxesenii* | chisel | kmr | NA | 1 | 35.87 | 0 | NA |
| 149 | gut | *Xi. saxesenii* | chisel | kmr | NA | 1 | 36.816 | 0 | NA |
| 149 | exterior | *Xi. saxesenii* | chisel | kmr | NA | 1 | 37.017 | 0 | NA |
| 150 | mycangia | *Xi. saxesenii* | chisel | kmr | NA | 0 | NA | 0 | NA |
| 150 | gut | *Xi. saxesenii* | chisel | kmr | NA | 1 | 36.119 | 0 | NA |
| 150 | exterior | *Xi. saxesenii* | chisel | kmr | NA | 1 | 36.691 | 0 | NA |
| 151 | mycangia | *Xi. saxesenii* | trap | ofr | dry | 1 | 36.282 | 0 | NA |
| 151 | gut | *Xi. saxesenii* | trap | ofr | dry | 1 | 36.3685 | 0 | NA |
| 151 | exterior | *Xi. saxesenii* | trap | ofr | dry | 1 | 38.262 | 0 | NA |
| 152 | mycangia | *Xi. saxesenii* | trap | kmr | flood | 0 | NA | 0 | NA |
| 152 | gut | *Xi. saxesenii* | trap | kmr | flood | 0 | NA | 0 | NA |
| 152 | exterior | *Xi. saxesenii* | trap | kmr | flood | 0 | NA | 0 | NA |
| 153 | mycangia | *Xi. saxesenii* | trap | kmr | flood | 0 | NA | 0 | NA |
| 153 | gut | *Xi. saxesenii* | trap | kmr | flood | 0 | NA | 0 | NA |
| 153 | exterior | *Xi. saxesenii* | trap | kmr | flood | 0 | NA | 0 | NA |
| 154 | mycangia | *Xi. saxesenii* | trap | kmr | dry | 0 | NA | 0 | NA |
| 154 | gut | *Xi. saxesenii* | trap | kmr | dry | 0 | NA | 1 | 35.009 |
| 154 | exterior | *Xi. saxesenii* | trap | kmr | dry | 0 | NA | 0 | NA |
| 155 | mycangia | *Xi. saxesenii* | trap | kmr | dry | 0 | NA | 0 | NA |
| 155 | gut | *Xi. saxesenii* | trap | kmr | dry | 0 | NA | 0 | NA |

| | | | | | | | | | |
|---|---|---|---|---|---|---|---|---|---|
| 155 | exterior | *Xi. saxesenii* | trap | kmr | dry | 0 | NA | 1 | 36.2195 |
| 156 | mycangia | *Xi. saxesenii* | trap | kmr | dry | 0 | NA | 1 | 35.724 |
| 156 | gut | *Xi. saxesenii* | trap | kmr | dry | 0 | NA | 1 | 36.397 |
| 156 | exterior | *Xi. saxesenii* | trap | kmr | dry | 0 | NA | 1 | 35.064 |
| 157 | mycangia | *Xi. saxesenii* | trap | kmr | dry | 0 | NA | 1 | 35.942 |
| 157 | gut | *Xi. saxesenii* | trap | kmr | dry | 0 | NA | 0 | NA |
| 157 | exterior | *Xi. saxesenii* | trap | kmr | dry | 0 | NA | 0 | NA |
| 158 | mycangia | *Xi. saxesenii* | trap | kmr | dry | 0 | NA | 1 | 36.149 |
| 158 | gut | *Xi. saxesenii* | trap | kmr | dry | 0 | NA | 1 | |
| 158 | exterior | *Xi. saxesenii* | trap | kmr | dry | 0 | NA | 0 | NA |
| 159 | mycangia | *Xi. saxesenii* | trap | kmr | dry | 0 | NA | 0 | NA |
| 159 | gut | *Xi. saxesenii* | trap | kmr | dry | 0 | NA | 0 | NA |
| 159 | exterior | *Xi. saxesenii* | trap | kmr | dry | 0 | NA | 1 | 35.829 |

**Table S3. ITS qPCR primer and probes for *Ceratocystis lukuohia* and *C. huliohia*.** The probes include a ZEN internal quencher and 6-FAM fluorophore. All primers and probes were designed by Heller et al. (2023).

| Target | Oligo name | Sequence (5' - 3') | Tm (˚C) |
|---|---|---|---|
| *C. lukuohia* | C.luku.ITS.For | CGTACCTATCTTGTAGTGAGATGAATGC | 62.3 |
| *C. lukuohia* | C.luku.ITS.Rev | GTTTACAGTGGCGAGACTTATATACTG | 61.3 |
| *C. huliohia* | C.huli.ITS.For | AAAACCTTATAGAAGGGGCCCCCCAACTAC | 62.3 |
| *C. huliohia* | C.huli.ITS.Rev | TTTTAGTGGTGAAGAAGATTACTTATACTG | 61.4 |
| *C. lukuohia* | C.luku.ITS.Pr | /56-FAM/CGGTRCCCT/ZEN/TCAGAAGGGCCCTACCAC/3IABkFQ/ | 71.5-73.6 |
| *C. huliohia* | C.huli.ITS.Pr | /56-FAM/AAAACCTTA/ZEN/TAGAAGGGGCCCCCCAACTAC/3IABkFQ/ | 69.8 |

**Table S4. Presence of *C. lukuohia* and *C. huliohia* detected through qPCR of the mycangia, gut, and exterior for each of the five ROD-associated beetle species.** The number positive (i.e., the number of samples in which the presence of *C. lukuohia* or *C. huliohia* was detected through qPCR) of the number assayed (i.e., the number of sampled body parts for that species) gives the percent positive (i.e., the percentage of samples positive for the presence of *C. lukuohia* or *C. huliohia*). The number assayed is not always consistent across the different body parts within one species because if the gut or mycangia of individual beetles was lost either in the chiseling or dissection process, the remaining body part of that individual was still processed. For the individual column, a positive individual refers to a beetle in which *C. lukuohia* or *C. huliohia* was detected in at least one of its body parts.

| Detection of *C. lukuohia* and *C. huliohia* | | | | | | | | |
|---|---|---|---|---|---|---|---|---|
| | Exterior | | Gut | | Mycangia | | Individual | |
| **Beetle species** | **(Number positive for *Ceratocystis* presence / Number assayed) % Positive** | | | | | | | |
| *X. affinis* | (5/9) | 55.6% | (7/9) | 77.8% | (6/8) | 75.0% | (9/9) | 100% |
| *X. ferrugineus* | (29/50) | 58.0% | (34/48) | 70.8% | (36/47) | 76.6% | (43/54) | 79.6% |
| *X. perforans* | (7/7) | 100% | (5/7) | 71.4% | (5/7) | 71.4% | (7/7) | 100% |
| *Xi. saxesenii* | (21/61) | 34.4% | (20/60) | 33.3% | (17/61) | 27.9% | (29/61) | 47.5% |
| *X. simillimus* | (6/32) | 18.8% | (17/29) | 58.6% | (7/23) | 30.4% | (21/32) | 65.6% |

**Table S5. Chi-squared tests and Fisher's exact tests assessing the difference in *C. lukuohia* and *C. huliohia* presence, respectively, amongst the body parts of each beetle species.** After Bonferroni corrections, significance level α = 0.00714. Significant p-values shown in bold.

| Beetle species | Pathogen | df | χ2 | p-value |
|---|---|---|---|---|
| *X. ferrugineus* | *C. lukuohia* | 2 | 6.2129 | 0.04476 |
| *X. ferrugineus* | *C. huliohia* | 2 | 1.3128 | 0.5187 |
| *Xi. saxesenii* | *C. lukuohia* | 2 | 0.84413 | 0.6557 |
| *Xi. saxesenii* | *C. huliohia* | 2 | 0.0841 | 0.9588 |
| *X. simillimus* | *C. lukuohia* | 2 | 10.918 | **0.004258** |
| *X. affinis* | *C. lukuohia* | 2 | *n/a (Fisher's test)* | 0.6567 |
| *X. perforans* | *C. lukuohia* | 2 | *n/a (Fisher's test)* | 0.4842 |

**Table S6. Pairwise comparisons from Generalized Linear Model (GLM) assessing the impact of beetle species, body part, and collection method on *Ceratocystis* presence in ROD-associated beetles.** Odds ratios (ORs) are the ratio of the odds of *Ceratocystis* presence in the category in the numerator to the odds of *Ceratocystis* presence in the category in the denominator. Bonferroni corrections were applied to all p-values. Significant p-values shown in bold.

| **Predictor (pairwise comparison)** | **OR**[1,2] | **95% CI**[2] | **p-value** |
|---|---|---|---|
| **Beetle species** | | | |
| *X. affinis* / *X. ferrugineus* | 3.01 | 0.74, 12.2 | 0.273 |
| *X. affinis* / *X. perforans* | 0.52 | 0.07, 3.75 | >0.999 |
| *X. affinis* / *Xi. saxesenii* | 5.08** | 1.41, 18.3 | **0.004** |
| *X. affinis* / *X. simillimus* | 13.0*** | 2.50, 67.3 | **<0.001** |
| *X. ferrugineus* / *X. perforans* | 0.17* | 0.03, 0.97 | **0.044** |
| *X. ferrugineus* / *Xi. saxesenii* | 1.69 | 0.73, 3.88 | 0.777 |
| *X. ferrugineus* /*X. simillimus* | 4.31*** | 1.74, 10.7 | **<0.001** |
| *X. perforans* / *Xi. saxesenii* | 9.72*** | 1.90, 49.7 | **<0.001** |
| *X. perforans* / *X. simillimus* | 24.8*** | 3.60, 171 | **<0.001** |
| *Xi. saxesenii* / *X. simillimus* | 2.55 | 0.78, 8.38 | 0.267 |
| **Body part** | | | |
| Exterior / Gut | 0.63 | 0.35, 1.12 | 0.165 |
| Exterior / Mycangia | 1.08 | 0.59, 1.96 | >0.999 |
| Gut / Mycangia | 1.72 | 0.95, 3.14 | 0.088 |
| **Collection method** | | | |
| Chisel / Trap | 3.13*** | 1.64, 5.97 | **<0.001** |

[1]*p<0.05; **p<0.01; ***p<0.001

[2]OR = Odds Ratio, CI = Confidence Interval

**Table S7. Pairwise comparisons from Generalized Linear Model (GLM) that assesses the impact of trap flooding on *Ceratocystis* presence in the beetles.** Beetle species, body part, and trap flooding are included as predictor variables. Only samples from beetles caught in traps were included in the model. Bonferroni corrections were applied to all p-values. Significant p-values shown in bold.

| **Predictor (pairwise comparison)** | **OR**[1,2] | **95% CI**[2] | **p-value** |
|---|---|---|---|
| **Beetle species** | | | |
| *X. affinis* / *X. ferrugineus* | 2.98 | 0.77, 11.6 | 0.203 |
| *X. affinis* / *X. perforans* | 0.53 | 0.08, 3.42 | >0.999 |
| *X. affinis* / *Xi. saxesenii* | 6.81*** | 1.81, 25.6 | **<0.001** |
| *X. affinis* / *X. simillimus* | 0.18* | 0.03, 0.94 | **0.036** |
| *X. ferrugineus* / *X. perforans* | 2.29 | 0.95, 5.53 | 0.079 |
| *X. ferrugineus* / *Xi. saxesenii* | 12.8*** | 2.51, 65.2 | **<0.001** |
| **Body part** | | | |
| Exterior / Gut | 1.20 | 0.56, 2.57 | >0.999 |
| Exterior / Mycangia | 1.24 | 0.58, 2.64 | >0.999 |
| Gut / Mycangia | 1.03 | 0.48, 2.22 | >0.999 |
| **Trap flooding** | | | |
| Dry / Flooded | 1.78 | 0.80, 3.97 | 0.158 |

[1]*p<0.05; **p<0.01; ***p<0.001
[2]OR = Odds Ratio, CI = Confidence Interval

**Table S8. Pairwise comparisons from Generalized Linear Mixed Model (GLMM) that includes the beetle collection site as a random effects variable to take into account collection site variability on *Ceratocystis* presence in the beetles.** Beetle species, body part, and collection method are included as fixed effects variables (referred to as "predictor variables" in GLMs). Bonferroni corrections were applied to all p-values. Significant p-values shown in bold.

| Predictor (pairwise comparison) | OR[1] | 95% CI | p-value |
|---|---|---|---|
| **Beetle species** | | | |
| *X. affinis* / *X. ferrugineus* | 3.48 | 0.84, 14.4 | 0.138 |
| *X. affinis* / *X. perforans* | 0.52 | 0.07, 3.76 | >0.999 |
| *X. affinis* / *Xi. saxesenii* | 9.21*** | 1.97, 42.9 | **<0.001** |
| *X. affinis* / *X. simillimus* | 49.0 | 0.79, 3,022 | 0.081 |
| *X. ferrugineus* / *X. perforans* | 0.15* | 0.03, 0.85 | **0.022** |
| *X. ferrugineus* / *Xi. saxesenii* | 2.64 | 0.91, 7.67 | 0.104 |
| *X. ferrugineus* /*X. simillimus* | 14.1 | 0.33, 595 | 0.476 |
| *X. perforans* / *Xi. saxesenii* | 17.7*** | 2.80, 112 | **<0.001** |
| *X. perforans* / *X. simillimus* | 94.1* | 1.35, 6,568 | **0.027** |
| *Xi. saxesenii* / *X. simillimus* | 5.32 | 0.18, 157 | >0.999 |
| **Body part** | | | |
| Exterior / Gut | 0.62 | 0.34, 1.12 | 0.153 |
| Exterior / Mycangia | 1.08 | 0.59, 1.98 | >0.999 |
| Gut / Mycangia | 1.75 | 0.95, 3.21 | 0.081 |
| **Collection method** | | | |
| Chisel / Trap | 3.75*** | 1.91, 7.35 | **<0.001** |

[1]*p<0.05; **p<0.01; ***p<0.001

[2]OR = Odds Ratio, CI = Confidence Interval
